# Single-cell profiling identifies clinically relevant interactions between tumor associated macrophages and blood endothelial cells in diffuse large B cell lymphoma

**DOI:** 10.1101/2022.12.08.519637

**Authors:** Juliette Ferrant, Simon Le Gallou, Francine Padonou, Simon Léonard, Ilénia Papa, Céline Pangault, Anne Barthel, Vincent Launay, Guillaume Manson, Bénédicte Hoareau, Laurent Deleurme, Céline Monvoisin, Benoit Albaud, Nathalie Van Acker, Camille Laurent, Francisco Llamas-Gutierrez, Henri-Alexandre Michaud, Nathalie Bonnefoy, Thierry Pécot, Karin Tarte, Mikael Roussel

**Affiliations:** Unité Mixte de Recherche UMR1236, Institut National de la Santé et de la Recherche Médicale, INSERM, Université Rennes 1, Etablissement Français du Sang Bretagne, Equipe labellisée Ligue, LabEx IGO, F-35000, Rennes, France; Centre Hospitalier Universitaire de Rennes, SITI, Pôle Biologie, F-35033, Rennes, France; Centre Hospitalier Universitaire de Rennes, Laboratoire d’Hématologie, Pôle Biologie, F-35033, Rennes, France; Centre Hospitalier de Bretagne Sud, Laboratoire de biologie médicale, F-56100, Lorient, France; Centre Hospitalier de Saint Brieuc, Service d’Hématologie, F-22000, Saint Brieuc, France; Centre Hospitalier Universitaire de Rennes, Service d’Hématologie, F-35033, Rennes, France; Sorbonne Université, Faculté de Médecine, UMS037, PASS, Plateforme de Cytométrie de la Pitié-Salpêtrière CyPS, F-75013 Paris; UMS Biosit, Flow Cytometry Facility H2P2, Université Rennes 1, INSERM, CNRS, F-35000, Rennes, France; Institut Curie, PSL University, ICGex Next-Generation Sequencing Platform, F-75005, Paris, France; Institut Curie, PSL University, Single Cell Initiative, F-75005, Paris, France; Centre Hospitalier Universitaire de Toulouse, Service d’Anatomie et Cytologie Pathologiques, Imag’IN Platform IUCT, et INSERM UMR1037, F-31059, Toulouse, France; Centre Hospitalier Universitaire de Rennes, Service d’Anatomie et Cytologie Pathologiques, F-35033, Rennes, France; IRCM, Université de Montpellier, INSERM, ICM, Plateforme de Cytométrie et d’Imagerie de Masse, F-34000, Montpellier, France; Univ Rennes, CNRS, INSERM, Biosit UAR 3480 US_S 01, F-35000 Rennes, France; Centre Hospitalier Universitaire de Rennes, Laboratoire d’Immunologie, Pôle Biologie, F-35033, Rennes, France

## Abstract

Diffuse large B cell lymphoma (DLBCL) and follicular lymphoma (FL) are the two most common B-cell lymphomas and are characterized by a dynamic crosstalk between tumor B cells and a heterogeneous tumor-supportive microenvironment, including immune, endothelial, and stromal components. Although their impact on the pathogenesis and prognosis of B-cell lymphoma has been acknowledged for years, tumor-associated macrophages (TAM) have not been extensively explored in DLBCL and FL. Herein, we investigate mononuclear phagocytes (MNP) heterogeneity at the single cell level and their potential co-regulation with the stromal and endothelial compartments in B-cell lymphoma lymph nodes compared to reactive secondary lymphoid organs, using a combination of mass cytometry, single cell RNA sequencing, and *in silico* approaches. We reveal a co-regulation between TAM and blood endothelial cells (BEC) in lymphoma. Moreover, we identify a specific interaction between Annexin A1 (ANXA1)-expressing BEC and formyl-peptide receptors (FPR1/2)-expressing monocytes/macrophages in DLBCL, which we confirm *in situ* by multiplex immunofluorescence and imaging mass cytometry. This crosstalk is associated to an immunosuppressive tumor microenvironment and an adverse prognosis in DLBCL.

## INTRODUCTION

Diffuse large B cell lymphoma (DLBCL) and follicular lymphoma (FL) are the two most common lymphomas, both deriving from germinal center (GC) B cells and characterized by aggressive versus indolent clinical course, respectively.^1^ Composition and organization of the immune and non-immune tumor microenvironment (TME) have been progressively identified as crucial in supporting FL and DLBCL tumor growth and resistance to treatment.^2–5^ Landscape of tumor-infiltrating T-cells and stromal cells has been explored at the single cell level in DLBCL and FL^6–9^ but the heterogeneity of the myeloid compartment still remains essentially uncharacterized, mainly due to the technical limitations of the classically used non-enzymatic lymph node (LN) dissociation approaches.

Conversely, the mononuclear phagocyte (MNP) compartment, encompassing macrophages, monocytes, and dendritic cells (DC), has hence been recently thoroughly explored in patients with solid cancers by single cell approaches, such as single cell RNA sequencing (scRNASeq) and mass cytometry (or cytometry by time-of-flight, CyTOF).^10^ More specifically, accumulation and phenotypic alterations of tumor-associated macrophages (TAM) have been associated with prognosis, resistance to therapy, and relapse in numerous solid tumors.^11,12^ TAM can display direct pro-tumoral activities and provide also indirect support to the tumor through a bi-directional crosstalk with other immune cells, essentially resulting in immunosuppression; and non-immune cells, including cancer-associated fibroblasts (CAF) and endothelial cells.^13^ In FL and DLBCL, several studies have shown an association between CD68^pos^ and/or CD163^pos^ macrophage infiltration and tumor prognosis, with sometimes contradictory results depending on the treatment schedule and likely the “low resolution” of the TAM phenotype, partly missing their functional heterogeneity.^14–20^ These discrepancies illustrate the ambivalent role of TAM, which can display pro-tumoral but also anti-tumoral properties in B-cell lymphomas, contributing in particular to the efficacy of immunochemotherapy based on anti-CD20 antibodies through antibody-dependent cell phagocytosis (ADCP). TAM display immunosuppressive functions in DLBCL via expression of PD-L1,^21–23^ and have low tumor phagocytosis activities in DLBCL and FL due to their overexpression of the phagocytosis-inhibitory receptor SIRPα.^24^ Furthermore, FL TAM directly promote FL B cell growth through trans-presentation of IL-15.^25^ FL TAM also express CD209/DC-SIGN which binds oligomannose residues occupying N-glycosylation sites introduced within variable regions of FL BCR (and less frequently DLBCL), leading to antigen-independent BCR-dependent B cell activation.^26–28^ FL stromal cells overexpressed CCL2, thus recruiting monocytes that further differentiate into pro-angiogenic and immunosuppressive TAM.^29^ The TAM pro-angiogenic properties were also demonstrated in a xenograft model of DLBCL.^30^ In normal lymph node (LN), highly specialised macrophages, stromal, and endothelial cells, involved at different levels in the immune response, interact within distinct niches.^31^ Recently, the crosstalk between blood vessel permeability, production of CCL2 by lymphoid stromal cells, and recruitment of monocytes into lymph nodes (LN) has been shown to regulate the behavior of normal B cells.^32^ However, the dialogue between TAM and stromal or endothelial cells in FL and DLBCL, and how it could impact lymphomagenesis has not been fully explored.

In the current study, we aimed at exploring the co-evolution of MNP, stromal, and endothelial heterogeneity within FL and DLBCL LN through a combination of mass cytometry, scRNAseq, and *in silico* approaches. We identified a coregulation between TAM and blood endothelial cell (BEC) activation status in lymphoma notably relying on the deregulation of the ANXA1/FPR1-2 axis. This predicted interaction was validated in situ by multiplex immunofluorescence (mIF) and imaging mass cytometry (IMC) and was associated with an adverse prognosis in DLBCL, identifying a new clinically relevant MNP-BEC loop in this disease.

## RESULTS

### Mononuclear phagocyte, stromal cell, and endothelial cell landscape differ across normal and B-cell lymphoma lymphoid tissues

MNP and stromal cells are often described as engaged in a bidirectional pro-tumoral crosstalk in solid tumors but are still poorly known in B-cell lymphoma. To fill this gap, we explored LN from FL and DLBCL at both the phenotypic and transcriptomic levels and compared them with their physiological counterpart reactive secondary lymphoid organs (rSLO), including reactive lymph nodes (rLN) and reactive tonsils (rTonsil) (Table 1 and Figure S1A). Firstly, we mechano-enzymatically dissociated tissue fragments from 24 SLO samples, including DLBCL (n=6), FL (n=9), rLN (n=3), and rTonsil (n=6), as previously described.^33^ Following T- and B-cell depletion, cells were stained with a dedicated CyTOF panel (Key resources table and Figure S1B). After pre-processing to exclude dead cells, doublets, and contaminating CD3^pos^ and CD19^pos^ cells, we obtained 594,946 CD45^neg^ viable cells (n = 79,955 for DLBCL, n = 105,344 for FL, n = 86,839 for rLN, and n = 322,808 for rTonsil) and 658,206 CD45^pos^ HLA-DR^pos^ MNP (n = 138,374 for DLBCL, n = 152,000 for FL, n = 236,902 for rLN, and n = 130,930 for rTonsil). CD45^neg^ and CD45^pos^ HLA-DR^pos^ cells were analyzed separately with the same pipeline including dimension reduction by UMAP, flowSOM clustering, and manual annotation of the clusters (Figure S1C).

**Table 1:** Patient characteristics.

| Disease/<br>sample<br>type | Tissue | Age at<br>diagnosis<br>(years) | Cell of origin<br>(DLBCL) or<br>grade (FL) | Treatment | CyTOF | scRNAseq | Hyperion |
| --- | --- | --- | --- | --- | --- | --- | --- |
| rTonsil | tonsil | 61 | NA | NA | X |  |  |
|  | tonsil | 40 | NA | NA | X | X |  |
|  | tonsil | 29 | NA | NA | X | X |  |
|  | tonsil | NA | NA | NA | X |  |  |
|  | tonsil | NA | NA | NA | X |  |  |
|  | tonsil | NA | NA | NA | X |  |  |
|  | tonsil | NA | NA | NA |  | X |  |
| rLN | LN | 60 | NA | NA | X | X | X |
|  | LN | 53 | NA | NA | X | X | X |
|  | LN | 19 | NA | NA | X | X | X |
| FL | LN | 85 | 1-2 | - | X | X | X |
|  | LN | 45 | 3a (relapse) | R-CHOP, Len | X | X |  |
|  | LN | 75 | 1-2 | - | X | X |  |
|  | LN | 65 | 1-2 (relapse) | R-CHOP | X | X |  |
|  | LN | 65 | 1-2 (relapse) | R-CHOP | X |  |  |
|  | LN | 79 | 1-2 | - | X |  | X |
|  | LN | 79 | 3a | - | X |  |  |
|  | LN | 67 | 1-2 (relapse) | R-CHOP | X |  |  |
|  | LN | 58 | 1 | - | X |  | X |
| DLBCL | LN | 74 | non-GC | - | X |  | X |
|  | LN | 69 | non-GC | - | X |  | X |
|  | LN | 52 | GC | - | X |  | X |
|  | LN | 74 | ABC | - | X | X | X |
|  | LN | 75 | non-GC | - | X | X |  |
|  | LN | 53 | GC | - | X | X |  |

\*replicate of same patient. ABC : Activated B cell, DLBCL : Diffuse Large B cell Lymphoma, FL : Follicular Lymphoma, GC : Germinal Center, Len : Lenalidomide, LN : Lymph Node, R-CHOP : Rituximab Cyclophosphamide Doxorubicine Vincristine Prednisone, rLN : reactive Lymph Node, scRNASeq : single cell RNASeq, NA : Not applicable.

CD45^pos^ HLA-DR^pos^ cells were separated in 13 clusters, including a CD123^pos^ pDC and 10 MNP clusters (Figure 1A). The 2 other clusters, named CD11b^hi^ cluster and CD209^hi^ cluster, comprised a small number of events (respectively 2,409 and 584 cells), were not identified as MNP, and were thus excluded from further analysis. Among the 10 MNP clusters we found i) 2 DC clusters (respectively named cDC [CCR2^lo^ SIRP*α*^hi^ HLA-DR^hi^ CD11c^dim^] and moDC [CCR2^pos^ SIRP*α*^hi^ HLA-DR^hi^ CD11c^pos^], ii) a CCR2^pos^ CD14^pos^ monocyte/macrophage (moMac) cluster, iii) a CCR2^pos^ CD14^neg^ CD11b^lo^ cluster that we could not identify as either monocyte/macrophage or monocyte/DC and thus named moMac/DC, and finally iv) 6 macrophage clusters (named Mac-CD169, Mac-CD209, Mac-PDL1, Mac-CD34, Mac-S100A9, and Mac-RANKL). Of note, the ratio between the sum of all monocyte and macrophage clusters (Mo&Mac metacluster) and the sum of DC clusters (DC metacluster) was enriched in lymphoma compared to rSLO (p < 0.001) (Figure 1B). Mac-CD209, Mac-CD34, Mac-S100A9, moMac, and moMac/DC were significantly enriched in lymphomas (p < 0.05) and almost absent in healthy tissues (Figures 1A and S1D).

**Figure 1:**
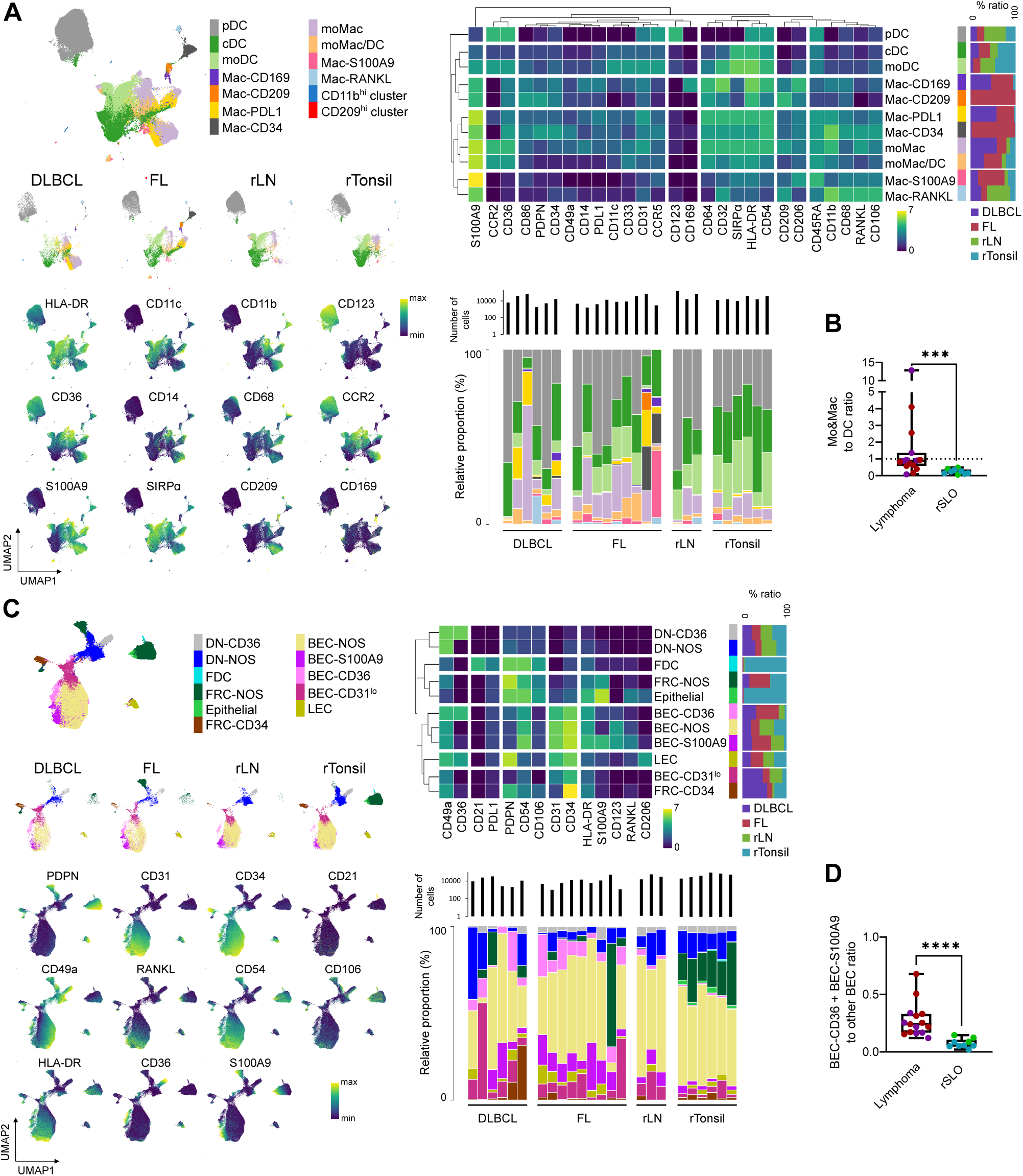
Stromal, endothelial, and MNP phenotypic landscape of reactive and lymphoma SLO. (A) UMAP and FlowSOM clustering of all 658,206 myeloid cells from SLO included in the CyTOF analysis and by sample (n = 138,374 DLBCL, n = 152,000 FL, n = 236,902 rLN, n = 130,930 rTonsil), displaying the clusters identified and UMAP projection of selected markers expression (left). Heatmap of the mean metal intensity (MMI) of selected markers and tissue abundance for the 11 MNP clusters, and relative proportion of the clusters for each sample (right). (B) Ratio of all monocytes/macrophages clusters (Mo&Mac metacluster) to all DC clusters, according to sample type. Mann-Whitney, ***p < 0.001. (C) UMAP and FlowSOM clustering of all 594,946 CD45^neg^ cells from SLO included in the CyTOF analysis and by sample type (n = 79,955 DLBCL, n = 105,344 FL, n = 86,839 rLN, n = 32,2808 rTonsil), displaying the clusters identified and UMAP projection of selected markers expression (left). Heatmap of the mean metal intensity (MMI) of selected markers and tissue abundance for the 11 stromal and endothelial clusters, and relative proportion of the clusters for each sample (right). (D) Ratio of BEC-CD36 and BEC-S100A9 clusters to other BEC clusters, according to sample type. Mann-Whitney, ****p < 0.0001. See also Figure S1.

Ten clusters of stromal and endothelial cells were identified within CD45^neg^ cells (Figure 1C). Among these 10 clusters, we found i) a CD31^neg^ PDPN^pos^ CD21^pos^ follicular dendritic cell (FDC) cluster, ii) 2 CD31^neg^ PDPN^pos^ CD49a^pos^ follicular reticular cell (FRC) clusters (FRC-NOS [not otherwise specified] and FRC-CD34 [that could also include pericytes and adventitial cells]), iii) 2 CD31^neg^ PDPN^neg^ CD49a^pos^ double negative (DN) clusters distinguished by CD36 expression (DN-NOS and DN-CD36), iv) a CD31^pos^ PDPN^pos^ lymphoid endothelial cell (LEC) cluster, and v) 4 CD31^pos^ CD34^pos^ PDPN^neg^ blood endothelial cell (BEC) clusters (BEC-NOS, BEC-CD36, BEC-S100A9, and BEC-CD31^lo^). Finally, a CD31^neg^ PDPN^pos^ CD49a^lo^ S100A9^pos^ cluster was present only in rTonsil and labelled as epithelial cells. Among BEC, BEC-CD36, and BEC-S100A9 were significantly enriched in lymphoma samples (p < 0.001) compared to rSLO (Figure 1D). Altogether this mapping of non-lymphocyte cells within SLO highlights co-amplification of specific subsets of macrophages and BEC in B-cell lymphomas.

### Mononuclear phagocytes and blood endothelial cells are co-regulated in SLO

We next evaluated whether MNP, stromal, and endothelial cells were co-regulated in SLO. To this aim, we combined on a hierarchical clustering both cluster abundances and mean metal intensities (MMI) (Figure S2A). Parameters showing statistical differences (Kruskal-Wallis p < 0.05) between sample types (FL, DLBCL, rLN, and rTonsil) (Figures S2A and 2B) were further used for sample re-clustering. Tonsil samples were clustered together and closely related to another cluster that include all rLN samples, 1 out of 9 FL and 1 out of 6 DLBCL samples (Figure 2A). A third cluster included exclusively FL samples (6 out of 9) and the last cluster included 5 DLBCL and 2 FL samples. rTonsils showed a high abundance of lymphoid stromal cells including FRC and FDC. Although rather heterogeneous, DLBCL samples displayed either i) an upregulation of CD54 on the DN-CD36 cluster, ii) an increase in the Mac-PDL1 percentage, and/or iii) an upregulation of CD36 on the BEC-CD36 cluster associated with an increase of FRC-CD34 and BEC-CD36 percentages. We noticed an upregulation of HLA-DR on the BEC clusters, also found in FL samples. In addition, FL samples were characterized by i) an upregulation of the TAM marker CD206 on several MNP clusters, ii) an increase of S100A9 or CD33/SIGLEC-3, involved in the inhibition of phagocytosis on MNP,^34^ iii) a high expression of CD31 and CD54 on BEC (Figure 2A). Altogether, BEC were activated in both FL and DLBCL compared to rSLO. To better evaluate how the parameters of interest could be co-regulated in SLO, we built a correlation matrix based on all LN samples (Figure 2B). Various BEC subsets, mostly overexpressing HLA-DR or CD31, were significantly co-regulated with CD33, S100A9, and CD206 expression on MNP subsets or with the abundance of Mac-S100A9 and Mac-PDL1. Altogether, these observations indicate that monocytes/macrophages and BEC are co-regulated in SLO with a co-amplification/co-activation in B-cell lymphoma compared to rSLO.

**Figure 2:**
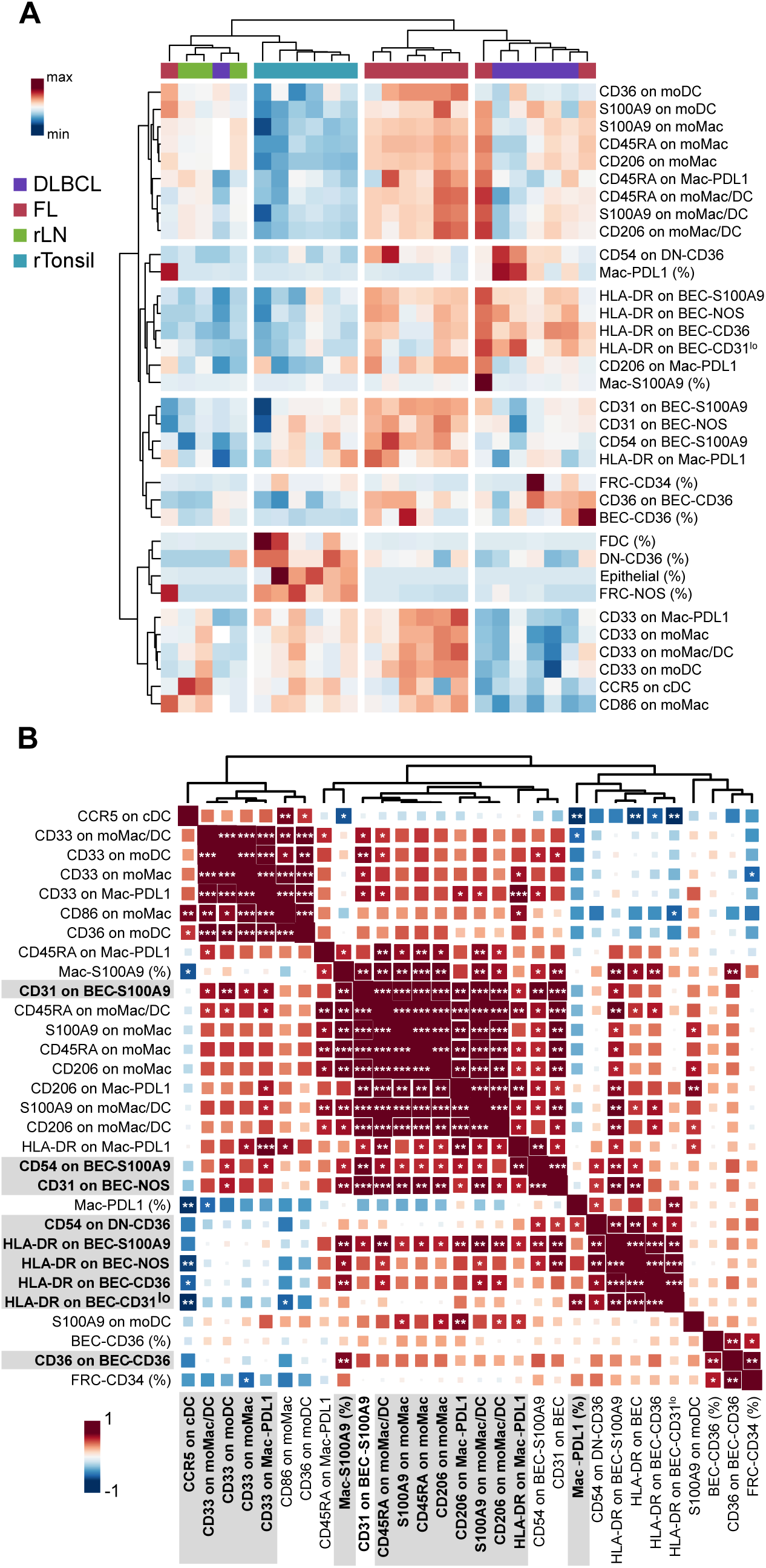
Myeloid subsets and Blood Endothelial Cells are co-regulated in SLO. (A) Heatmap of the statistically different parameters (abundance of clusters and mean metal intensity between sample types (FL, DLBCL, rLN, rTonsil). (B) Correlation matrix displaying the same parameters (except FDC, FRC-NOS, Epithelial cells, and DN-CD36 abundances) for LN samples (rLN, FL, and DLBCL). Spearman test, *p<0.05, **p<0.01, ***p<0.001. Parameters involved in significant correlations between myeloid and stromal/endothelial populations are highlighted in grey. See also Figure S2.

### Mononuclear phagocyte landscape in reactive SLO and lymphoma as revealed by scRNASeq

We then used scRNAseq to better explore lymphoma MNP heterogeneity from 13 samples (3 DLBCL, 4 FL, 3 rLN, and 3 rTonsil) at the transcriptomic level (Table 1). After sample preparation (mechano-enzymatic dissociation followed by T- and B-cell depletion), MNP were FACS-sorted (Figure S1A). All samples were already included in the CyTOF analysis except for one rTonsil (Table 1). We sorted and pooled CD45^pos^ CD3^neg^ CD19^neg^ HLA-DR^pos^ CD11c^pos^ and CD45^pos^ CD3^neg^ CD19^neg^ HLA-DR^pos^ CD11c^neg^ CD123^neg^ cells in order to eliminate pDC without losing any MNP subset (Figure S3A). A CITE-Seq analysis using CD14 and CD36 antibodies was used to better discriminate monocytes and macrophages from DC. After filtering and integration (Figure S3B), we obtained 13,229 cells (n = 2,225 DLBCL, n = 2,436 FL, n = 3,769 rLN, and n = 4,799 rTonsil) grouped into 17 clusters (Figure S3C). Among these 17 clusters, we selected the 7 clusters of MNP (moMac, Mac-C1Q, SIGLEC6-cDC, CLEC9A-cDC1, LAMP3-cDC, cDC2, MNDA-cDC2, altogether: 8,691 cells [n = 1,154 DLBCL, n = 1,444 FL, n = 3,091 rLN, and n = 3,002 rTonsil]) and performed a new integration, dimension reduction by UMAP, followed by a clustering (Figure 3A). Twelve clusters were identified and annotated, including one consisting of dying cells (117 cells, enriched in mitochondrial genes) excluded from further analysis. We identified i) 6 clusters of cDC including CLEC9A-cDC1, LAMP3-cDC, SIGLEC6-cDC (these last 2 subsets of cDC being recently described in several scRNASeq studies)^35–40^ and 3 subsets of CLEC10A^pos^ CD1C^pos^ cDC2 (MNDA-cDC2, CD1E-cD2, and CREM-cDC2), ii) 1 cluster of macrophages (Mac-C1Q), and iii) 4 monocyte/macrophage clusters (CD14-moMac, S100-moMac, CD16-moMac, and IL1b-moMac) (Figures 3B and 3C). Similar to what we observed in the CyTOF data, the ratio between Mo&Mac and DC metaclusters was increased in lymphoma compared to rSLO (Figure 3D). CD14-moMac and S100-moMac were *CD16*^lo^ *SOD2*^hi^ and were distinguishable from each other by the expression of *S100A8, S100A9, VCAN*, and *CEBPB*. Since S100-moMac showed high expression of *S100A8/9, VCAN,* and *CEBPB*, they could correspond to MDSC/TAM-like monocytes (Figures 3B and 3C).^10,41^ CD16-moMac were CD14^neg^ and expressed *FCGR3A/CD16* as well as high levels of *MAFB* and *CD68*, and are likely non-classical monocytes (ncMo)-like Mo&Mac. The IL1b-moMac cluster displayed high levels of the pro-inflammatory *IL1B* gene, as well as *FOS* and *JUN* that are involved in many cellular processes including cell differentiation, and could be inflammation-driven. Finally, only the Mac-C1Q macrophage cluster was significantly enriched in DLBCL (Figures 3E and S3D) and expressed high levels of *CD68*, of the 3 subunits of the complement subcomponent C1q: *C1QA, C1QB*, and *C1QC*, and of the matrix metalloproteinase *MMP9,* involved in extracellular matrix remodelling, angiogenesis, and neovascularization.

**Figure 3:**
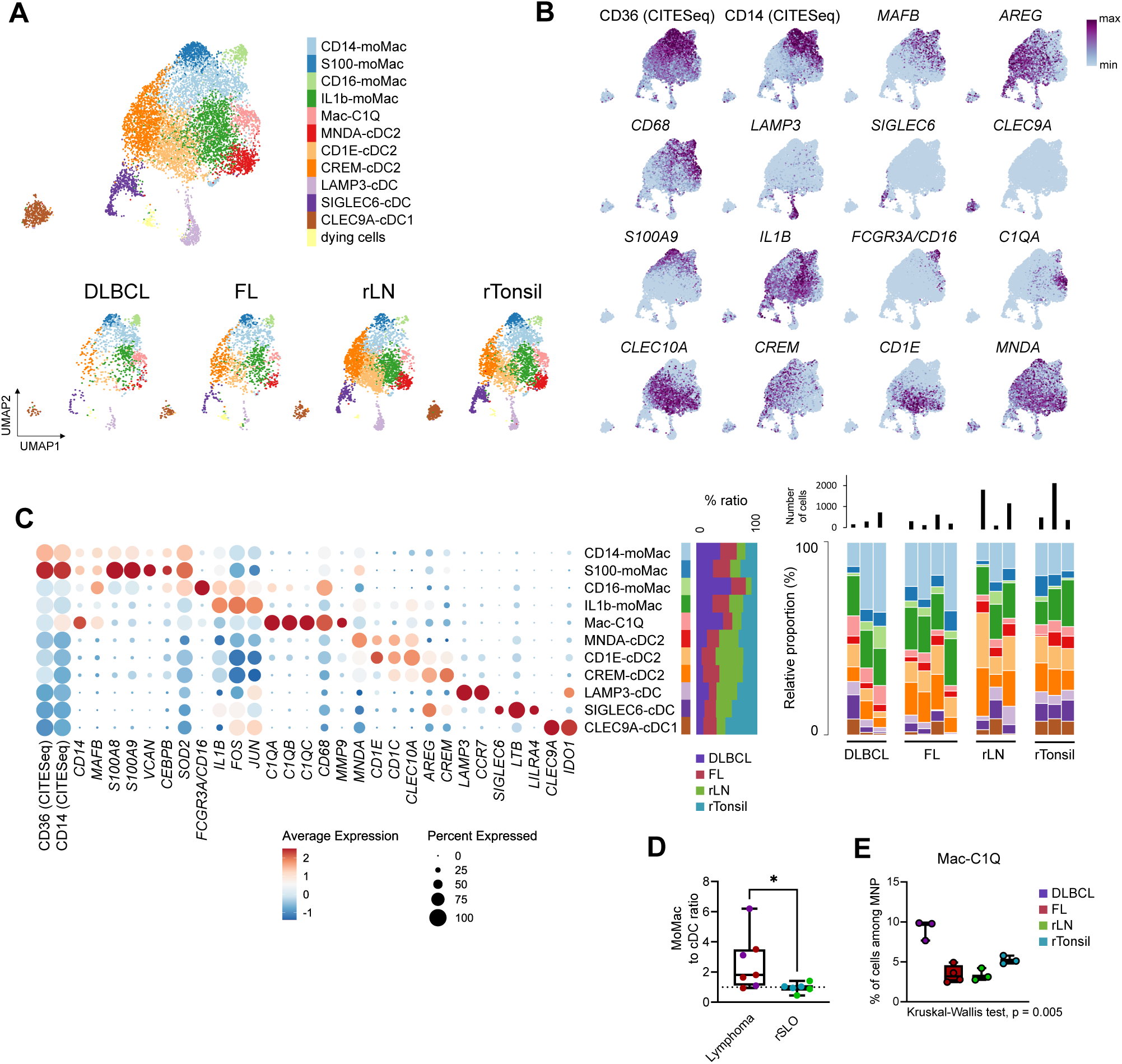
MNP landscape in reactive and lymphoma SLO as revealed by scRNASeq. (A) UMAP and FlowSOM clustering of all 8691 cells from SLO included in the single cell RNASeq analysis and by sample (n = 1154 DLBCL, n = 1444 FL, n = 3091 rLN, n = 3002 rTonsil), displaying the 12 clusters identified, including 11 MNP clusters. (B) UMAP projection of selected markers expression. (C) Dotplot showing the average RNA expression level (color scale) and percentage of expressing cells (dot size) of selected markers for the 11 MNP clusters (left), tissue preference of each cluster (center), relative proportion of the clusters for each sample (right). (D) Ratio of all monocytes/macrophages clusters (Mo&Mac metacluster) to all cDC clusters, according to sample type. Mann-Whitney, *p < 0.05. (E) Boxplot of Mac-C1Q cluster abundance according to sample type. See also Figure S3.

### Monocyte and macrophage deregulation at the transcriptomic level is disease specific

Due to the low abundance of cDC in the lymphoma samples and our previous results showing a correlation between monocytes/macrophages and BEC, we then focused on the transcriptomic deregulation in the cells included in the Mo&Mac metacluster (including CD14-moMac, S100-moMac, CD16-moMac, IL1b-moMac, Mac-C1Q) (Figure 3C). To avoid bias in the differential expression analysis due to the limited number of samples, we performed iterative analyses with all samples but one and retained only the 651 (out of 1,937) genes statistically differentially expressed after this bootstrap analysis. We then filtered 393 genes of interest based on a fold change |FC|>1.5 and an expression by more than 30% of cells in at least one of the conditions tested (Table S1). As reported for stromal cells, Mo&Mac transcriptomic signature was different between rTonsils and rLN (Figure S4A). We thus focused our analysis on lymphomas and rLN samples. DLBCL was characterized by a high number of genes differently expressed when compared to FL and rLN (respectively 174 and 268 genes), whereas less genes distinguished FL and rLN (33 genes) (Figure 4A). We defined a signature of 73 upregulated genes specific of Mo&Mac from DLBCL, which includes i) hallmark macrophage/TAM genes (*CD14, CD163, FCGR1A/B*) and non-classical monocyte marker (*FCGR3A/CD16*), i) gene involved in immune suppression (*IL4I1*), and iii) genes from the S100 family (*S100A8, S100A9*) and interferon (*IFN*) response genes (*IFITM1, IFITM3, IFI30, STAT1*, and the chemokines *CXCL9* and *CXCL10*) (Table S1). Other genes were found upregulated in macrophages from DLBCL vs either FL (26 genes, including *C1QB/C*) or rLN (47 genes, including *S100A4/6/11/12* from the S100 family, the neutrophil-attractant chemokine *CXCL8*, the macrophage transcription factor *MAFB*, and the macrophage/TAM marker *VCAN*). We also performed the differential expression analyses separately on each cluster previously described (Table S2). No gene was deregulated in the CD16-moMac subset, which could be linked to the low number of cells. Altogether, Mo&Mac clusters from DLBCL were enriched in genes involved in immune suppression (*IDO1, CD274*) and in IFN signalling (*IFI6, TRIM22, IFITM2*) (Table S2). Genes downregulated in FL and DLBCL compared to rLN were often hallmark DC genes and/or involved in antigen presentation (including *CD74* and *MHCII* genes such as *HLA-DRB1, HLA-DQA1, HLA-DPB1*). Only 2 genes (*CALHM6* and *IFITM3*) were specifically downregulated in FL, both involved in response to IFN, possibly reflecting a relative non-inflammatory environment in FL, at least compared to the reactive and aggressive lymphoma settings. Among the 14 upregulated genes in FL compared to rLN, we found 3 genes from the S100 family (*S100A6/8/9*) and the M-ficolin gene (*FCN1*). We then performed an analysis of the deregulated pathways between DLBCL, FL, and rLN, with g:profiler, followed by an enrichment map analysis with Cytoscape^42^ (Figure 4B). Several pathways were upregulated in DLBCL (highlighted in purple) and were involved in chemotaxis, pattern recognition receptor (PRR) signalling, immune response regulation, and IFN signalling pathway. Another group of pathways (highlighted in yellow) involved in translation was downregulated in DLBCL compared to the other two conditions. Finally, two clusters of pathways were downregulated in both DLBCL and FL compared to rLN, involved in antigen presentation to T cell via MHC class II (highlighted in green).

**Figure 4:**
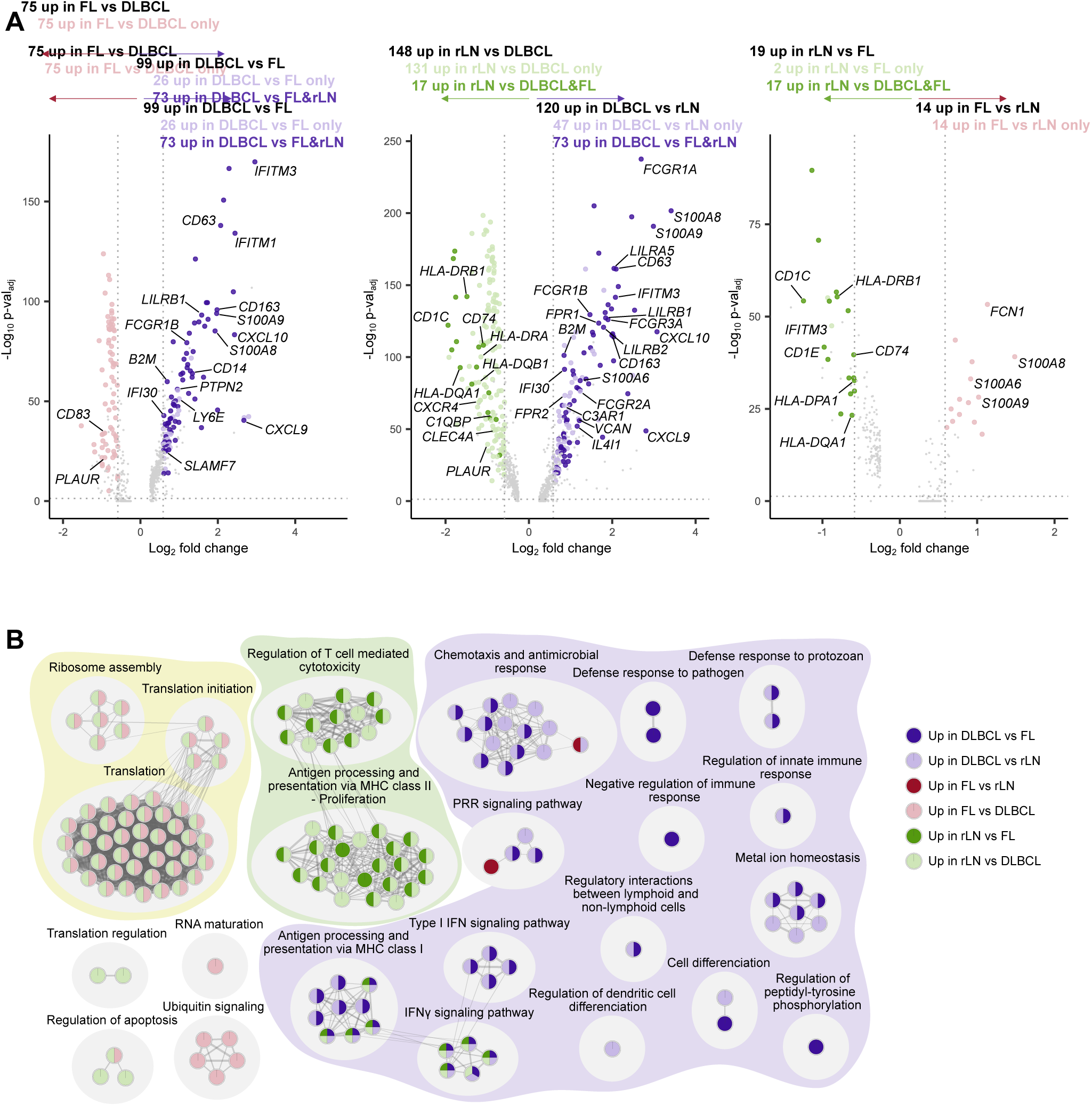
Monocytes and macrophages transcriptomic deregulation in lymphoma. (A) Volcano plots of differentially expressed genes for monocytes and macrophages included in the CD14-moMac, S100-moMac, CD16-moMac, IL1b_moMac/DC, and Mac-C1Q, between FL and DLBCL (left), DLBCL and reactive LN (center), FL and reactive LN (right). Wilcoxon-Mann-Whitney test, |FC| > 1.5, p<0.05. Only genes present in at least 30% of the cells in one of the compared conditions and found to be robustly differentially expressed by iteratively removing each sample from the differential analysis are presented as differentially expressed (larger, colored dots). Genes found to be deregulated in the comparison between clusters of one sample type and the other two sample types are highlighted in darker colors. (B) Enrichment map of the pathway analysis for the differentially expressed genes. Each node represents a pathway, colored according to the context in which it is enriched. See also Figure S4.

Then, we used the PISCES R package^43^ on our scRNAseq dataset to infer protein activity. As previously done with the differentially expressed genes, we filtered significantly dysregulated inferred proteins, either in Mo&Mac metacluster (Table S3) or in each individual cluster (Table S4). We observed a strong DLBCL Mo&Mac signature including macrophage/TAM markers (MAFB, PPARG) and IFN response (STAT1, STAT2, IRF1, IRF9) (Figures S4B and S4C). The MDSC marker STAT3 was among the few inferred proteins upregulated in FL compared to rLN, as well as in DLBCL vs rLN. Overall, monocyte and macrophage transcriptome shows a significant deregulation in DLBCL, compared to both rLN and FL, whereas FL MNP niche shares features with both reactive and DLBCL LN.

### Interactome and *in situ* analyses identify BEC and Mo&Mac crosstalk in DLBCL

We next sought to extend our previous observations of co-amplification of macrophage and BEC subsets in B-cell lymphomas, by predicting *in silico* potential receptor-ligand interactions. CellPhoneDB was used to predict receptor-ligand interactions between macrophages and BEC in normal vs lymphoma (DLBCL or FL) conditions.^44^ We first defined Mo&Mac genes of interest by combining genes previously defined in Tables S1, S2, S3, and S4 (i.e. genes that were up- or down-regulated either in DLBCL or in FL compared to rLN, and obtained by scRNAseq or corresponding to the inferred proteins; both either from Mo&Mac metacluster or from one of the clusters). We took advantage of already published scRNASeq data of BEC from the Eµ-*Myc* mouse model of aggressive lymphoma compared to control mice.^45^ Seven statistically significant interactions were identified between ligands expressed by BEC and receptors expressed by Mo&Mac, in either the lymphoma context, the reactive context, or both (Figure 5A). Lymphoma BEC were predicted to interact with Mo&Mac from DLBCL via AnnexinA1 (ANXA1) binding to the formyl-peptide receptors FPR1 or FPR2, an interaction not found in reactive context. Accordingly, ANXA1 was upregulated in lymphoma BEC (Figure 5A) whereas FPR1 and FPR2 were found upregulated in DLBCL Mo&Mac metacluster and in most subclusters including the Mac-C1Q and CD14-moMac clusters (Figures 5A, 5B, and S5A). Conversely, CXCL12-CXCR4 and ICAM1-AREG interactions seemed to be lost in DLBCL as compared to reactive context (Figure 5A). No predicted lymphoma-specific interaction was found in the reverse analysis (i.e. ligands expressed on Mo&Mac and receptors on BEC) (Figure S5B). Regarding the Mo&Mac genes deregulated in FL, no specific interaction was found between BEC and Mo&Mac (data not shown). We confirmed the putative interaction between BEC and Mo&Mac metacluster in DLBCL via ANXA1 binding to FPR1 or FPR2 by using the NicheNet algorithm^46^ (Figure S5C). On bulk RNASeq in Eµ-*Myc* vs control mice,^47^ *ANXA1* was overexpressed only by BEC and not by LEC, FRC, and DN, whereas *FPR1* was not overexpressed on these cells and FPR2 was overexpressed on BEC and DN (Figure S5D). *ANXA1*, *FPR1*, and *FPR2* were not differentially expressed by T and B cells in a scRNASeq dataset from human DLBCL and rLN samples^9^ (Figure S5E), and *ANXA1* was not differentially expressed by Mo&Mac in our own scRNASeq dataset (Figure S5F), therefore supporting ANXA1-FPR1/FPR2 interaction as a specific actor of BEC and Mo&Mac crosstalk in DLBCL. We confirmed this interaction *in situ* with FPR1-expressing CD68^pos^ macrophages found in close contact with ANXA1-expressing CD31^pos^ endothelial cells in DLBCL (Figure 5C), a pattern not found in rLN (Figure S5G).

**Figure 5:**
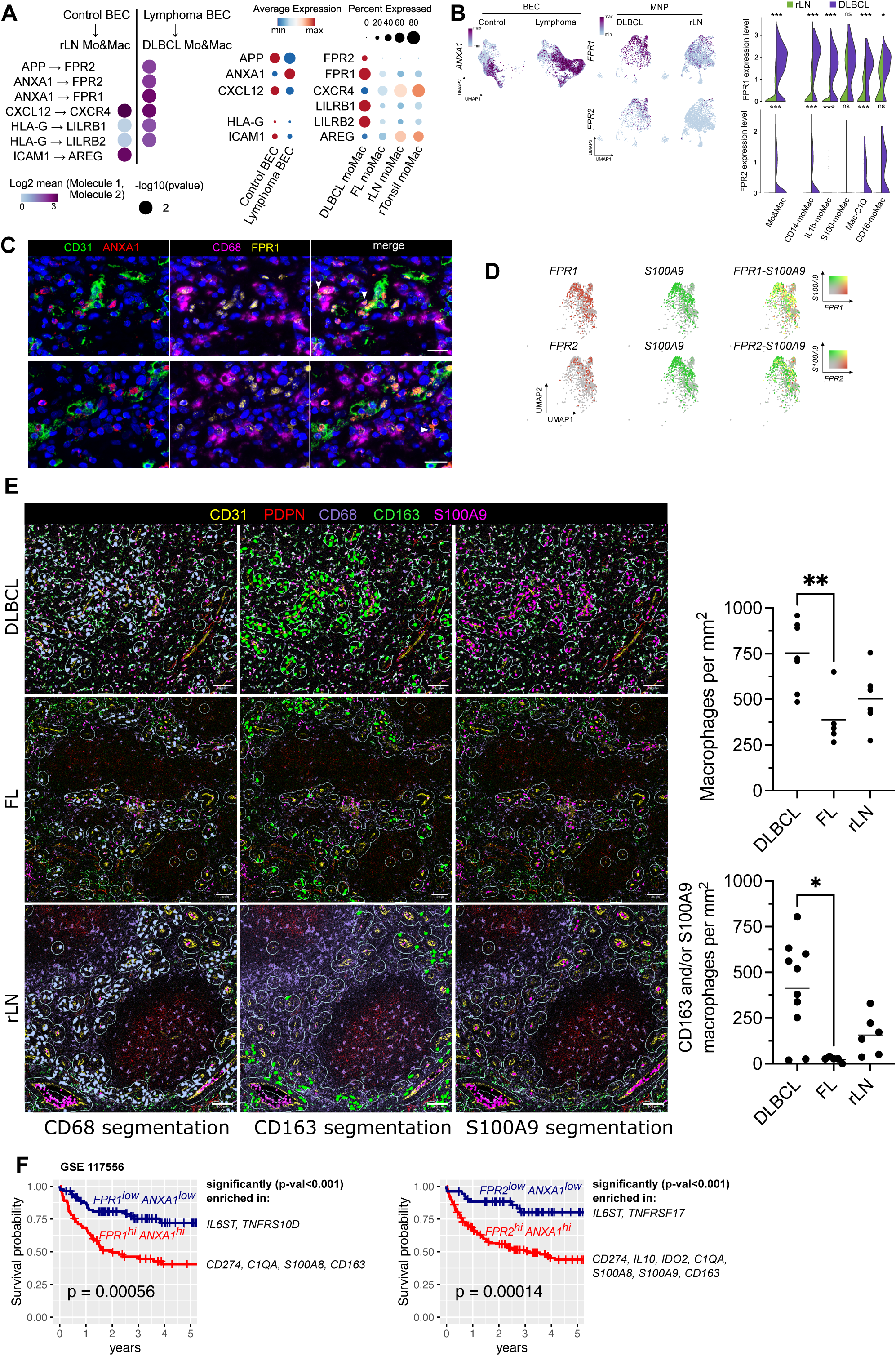
Interactome and *in situ* analyses identifies BEC and Mo&Mac crosstalk in DLBCL. (A) Dotplot of statistically significant interactions identified with CellphoneDB between normal murine BEC and rLN Mo&Mac or tumor murine BEC and DLBCL Mo&Mac, involving ligands expressed by murine BEC and receptors expressed by Mo&Mac and differentially expressed between rLN and DLBCL LN. Log2 mean expression of interacting molecules (color scale) and p-values (dot size) are shown. Only statistically significant interactions are plotted (p < 0.05) with a minimal p-value set to 0.01 (left). Dotplots showing the average RNA expression level (color scale) and percentage of expressing cells (dot size) of the identified ligands expressed by tumor or normal BEC and receptors expressed by Mo&Mac (right). (B) *ANXA1* expression on normal and tumor BEC^45^ and *FPR1* and *FPR2* expression levels on DLBCL Mo&Mac clusters (left). Mann-Whitney, *p < 0.05. (C) CD31, ANXA1, CD68, and FPR1 staining by mIF in DLBCL. Arrows indicate colocalization. Scale bar represent 20 µm. (D) *FPR1, FPR2* and *S100A9* expression levels on DLBCL Mo&Mac clusters. (E) IMC staining for BEC (CD31^pos^ PDPN^neg^), LEC (CD31^pos^ PDPN^pos^), and macrophages/myeloid cells (CD68, CD163, and S100A9) in 25 uM vicinity area of blood vessels. ROI for DLBCL (n=4, 10 ROI, total area 7.1 mm^2^), FL (n=3, 5 ROI, total area 5.6 mm^2^), and rLN (n=3, 6 ROI, total area 6.5 mm^2^). Blood vessels were defined as anatomical structure CD31^pos^ PDPN^neg^. CD31 cells are in yellow; PDPN are in red, CD68 are in purple, CD163 are in green, and S100A9 are in pink; neighbourhoods’ zones of 25 µm around blood vessels are indicated in light green. Representative ROI for DLBCL, FL, and rLN. Number of macrophages and macrophages expressing CD163 and/or S100A9 in 25 µm vicinity area of blood vessels. Kruskal-Wallis with Dunn’s multiple comparisons correction, *p < 0.05. Scale bar represent 100 µm. (F) Kaplan-Meier survival curves and log-rank test comparing patients with high proportion of *FPR1* and *ANXA1 (top)* and high proportion of *FPR2* and *ANXA1 (bottom)* to those with low proportion (GSE117556).^48^ Selected genes statistically enriched in groups of patients are indicated. See also Figures S5 and S6.

To confirm the spatial relationship between macrophages and BEC in DLBCL, we performed an imaging mass cytometry (IMC) analysis on DLBCL (n=4; 10 ROI [Region Of Interest]; total area 7.1 mm^2^), FL (n=3; 5 ROI; total area 5.6 mm^2^), and rLN (n=3; 6 ROI; total area 6.5 mm^2^) (Table 1). In our scRNAseq data, *S100A9* was among the top 10 genes correlated with both *FPR1* and *FPR2* (Table S5, Figure S5H) and FPR1 and FPR2 were coexpressed with S100A9 (Figure 5D), making S100A9 a good surrogate marker of FPR1/FPR2-expressing macrophages in DLBCL. Blood vessels were defined as CD31^pos^ PDPN^neg^ anatomical structures (Figures 5E and S6). On whole ROI, blood vessel area was higher in rLN and both CD68^pos^ and CD163^pos^ macrophages were enriched in DLBCL (p < 0.01) (Figure S6). To evaluate BEC/Mo&Mac interaction, we defined neighbourhood zones of 25 µm around the blood vessels and quantified CD68, CD163, and S100A9 cells located in this vicinity area (Figures 5E). Whole CD68^pos^ macrophages (p < 0.01) but also macrophage subsets expressing S100A9 and/or CD163 (p < 0.05) were enriched in the 25 µm area of blood vessels in DLBCL compared to FL.

Finally, we analysed the clinical impact of *ANXA1*, *FPR1,* and *FPR2* expression in a cohort of DLBCL samples.^48^ Log-rank test showed a statistically significant association between high *FPR1* and *ANXA1,* and high *FPR2* and *ANXA1* expression and a shorter overall survival (p = 5.6 x10^-4^ and p = 1.4 x10^-4^, respectively) (Figure 5F). Patients *FPR1*^hi^ and *ANXA1*^hi^ or *FPR2*^hi^ and *ANXA1*^hi^ showed an enrichment in immunosuppressive genes (*CD274, IL10, IDO2, C1QA, S100A8, S100A9, CD163*). These results suggest that Mo&Mac and BEC engage in a specific interaction involving the receptors FPR1 and FPR2 and their ligand ANXA1 in DLBCL, an interaction that may contribute to local immunosuppression and be linked to the prognosis of this aggressive lymphoma.

## DISCUSSION

MNP play pleiotropic roles in cancer pathogenesis, prognosis, and treatment. Their heterogeneity has been recently unraveled in large scRNAseq studies including numerous tumor tissues.^10,36,49^ However, these human tumor MNP atlases include very few lymphoma-derived cells obtained from poorly characterized lymphoma samples, precluding any conclusion on the lymphoma MNP landscape. Conversely, scRNAseq studies designed to specifically explore malignant B cells and their TME in FL or DLBCL do not include MNP, due to the use of purely mechanical dissociation approaches.^6,8,9^ In the current work, using a validated mechano-enzymatic dissociation and gating strategy,^33^ we provided a thorough characterization of human MNP in normal and malignant SLO making it possible to highlight their phenotypic and functional heterogeneity as well as their capacity to interact with other cell components of the TME, in particular BEC.

This approach allowed us to identify several Mo&Mac populations similar to those previously described in solid cancers, including CD14-moMac and CD16-moMac, resembling classical and non-classical tumor-infiltrating monocytes, and Mac-C1Q, previously proposed as long-term resident macrophages enriched in liver tumors.^36^ Our IL1b-moMac cluster displayed a profile of the previously described inflammatory cytokine-enriched TAM subset. Finally, S100-moMac expressed high amounts of *VCAN,* and *CREBBP*, but also *VEGFA*, and were previously identified as pro-angiogenic TAM subsets.^10^ Comparison between lymphoma and rSLO highlighted that DLBCL Mo&Mac are over-represented in tumor tissues, showing a specific transcriptomic profile, associated with an increase of immunosuppressive genes, including *CD274/PD-L1*, *IDO1*, and *LILRB1/LILRB2*, interferon-stimulated genes (ISG), and S100 genes. An immunosuppressive phenotype of DLBCL TAM has already been proposed, characterized by an upregulation of PD-L1, identified as a prognostic marker of the disease.^22,23,50^ Similarly, IDO1 expression and activity were also correlated with DLBCL patient outcome.^51,52^ Finally, LILRB1/2 have been identified as “don’t eat me” signals after engagement by MHC class I and could thus contribute to impairment of apoptosis previously reported in DLBCL.^24,53^ Interestingly, *LILRB2* and *IDO1* were found coregulated with PD-L1 expression on CD68^pos^ TAM in DLBCL contributing to a poor prognosis signature.^54^ Overall, analysis of large datasets of bulk RNAseq recently underlines the pro-tumoral role of TAM, revealing the co-occurrence of a high amount of TAM, a proinflammatory gene expression profile, and an overexpression of *PD-L1, IDO1*, and immunosuppressive cytokines.^3^ Conversely, the relationship between TAM and angiogenesis in B-cell lymphoma is largely underexplored even if a correlation between CD163^pos^ TAM and neovascularization through angiogenic sprouting has been proposed in FL.^55^

We showed here specific co-amplification and co-activation of TAM and BEC subpopulations in B-cell lymphomas compared to rSLO. In particular, suppressive PD-L1^pos^ TAM were correlated to HLA-DR expression on BEC in DLBCL, while upregulation of CD206 and S100A9 on different TAM subsets was correlated to an upregulation of CD31 and CD54 on BEC subtypes in FL. An increased density and disorganization of the vascular network is known in DLBCL,^56,57^ and neoangiogenesis signature is associated with adverse prognosis in this disease.^48^ In FL LN, although micro-vessel density seems to be equivalent to that of the reactive LN, specific angiogenic pathways are activated in endothelial cells.^7,56^ We identified an upregulation of HLA-DR on the BEC clusters in FL and DLBCL, as well as a significant enrichment of BEC-CD36 and BEC-S100A9, which probably reflects endothelial activation. Notably, CD36 has been directly involved in the neoangiogenesis process.^58,59^ Exploring the mechanism underlying the BEC/TAM crosstalk, we combined our MNP scRNASeq data with published murine BEC scRNASeq data, and predicted a specific paracrine interaction between ANXA1-expressing BEC and FPR1/FPR2-expressing Mo&Mac in DLBCL, with a possible additional autocrine ANXA1-FPR2 signalling in BEC. Of note, *ANXA1, FPR1,* and *FPR2* were not differentially expressed in FL samples compared to rLN in published datasets of stromal, endothelial, T, and B cells.^7,9^ ANXA1 is an anti-inflammatory calcium-regulated protein, whose only know receptors are the FPR G-protein coupled PRR formyl-peptide receptors. ANXA1-FPR interaction is involved in monocyte recruitment and polarization, switching them to an M2/reparative/pro-tumor phenotype.^60–64^ ANXA1 and/or FPR1/2 are dysregulated in several solid tumors, with a correlation to poor prognosis. The involvement of ANXA1-FPR signalling in tumor progression and tumor cell invasion has been mostly explored in autocrine tumor cell activation, notably in breast cancer, gastric cancer, pancreatic carcinoma, and glioblastoma.^65–68^ However, it was also involved in paracrine interactions between metastatic mammary cancer cells and microglia, driving microglia recruitment and polarisation to a tumor-supportive phenotype.^69^ Accordingly, elevated ANXA1, FPR1, and FPR2 are associated to poor outcome in breast cancer patients.^69^ Similarly, ANXA1-FPR1 interaction has been predicted in multiple myeloma between inflammatory mesenchymal stromal cells and myeloid cells, thus contributing to the formation of the protumoral bone marrow niche.^70^ Finally, cleaved ANXA1 has been described in caveolae of endothelial cells in several solid tumor tissues, specifically in the tumour vasculature, and not in the normal vessels.^65^ FPR1/2 expression was not cluster-specific in DLBCL Mo&Mac, and S100A9 was a surrogate marker for FPR1/2, allowing us to confirm in situ the BEC-TAM interaction. S100A9 is highly expressed in circulating monocytes/M-MDSC in DLBCL and contributes to their immunosuppressive activity.^71,72^ Moreover, DLBCL TAM overexpress CCR2, which could reflect a recent recruitment to the tissue.^50^ We also found a global upregulation of the monocyte markers CD14 and CD16 in Mo&Mac DLBCL compared to FL and rLN and CD16 was highly correlated to FPR1/2 expression in DLBCL Mo&Mac. An increased number of circulating CD16^pos^ ncMo, which display a combined inflammatory/tolerogenic phenotype and can be recruited by tumor B cells, has been shown in DLBCL.^73^ All these observations suggest an increased recruitment of monocytes, including ncMo, to the tumor in DLBCL, in which the ANXA1-FPR interaction could be a key player in the monocyte-derived TAM infiltration.

*ANXA1*, *FPR1,* and *FPR2* expression were shown to correlate to patient prognosis and expression of immunosuppressive genes in DLBCL samples. Neoangiogenesis and infiltration of immunosuppressive TAM subsets have been previously associated to poor prognosis in DLBCL. Our data suggest that both processes could be linked, through an ANXA1-FPR1/2 interaction involving BEC and Mo&Mac, leading to an infiltration of immunosuppressive TAM. Targeting TAM recruitment, survival, and function has become a major therapeutic goal in lymphoma and a specific focus on ANXA1-FPR1/2-mediated interactions may be of clinical interest in DLBCL.

### Limitation of the study

Although combining phenotypic and transcriptomic approaches, our single cell analyses are based on a limited number of samples. DLBCL is a clinically and biologically heterogeneous disease and the more recent DLBCL classifications reveal a relationship among tumor cell-of-origin, genomic alterations, and TME composition, including TAM content.^3,74^ Assessment of ANXA1 and FPR1/2 expression by BEC and Mo&Mac in larger populations of DLBCL patients would allow the identification of a potential association between ANXA1-FPR interaction and peculiar DLBCL molecular and TME subtypes. Moreover, we cannot exclude a loss of some macrophages during single cell suspension preparations, especially since macrophages are large, sometimes polynucleated cells embedded in the extracellular matrix that may be fragmented during isolation.^75^ Additionally, we used murine BEC data for in silico identification of BEC-Mo&Mac interactions. Finally, other co-regulations and crosstalk mechanisms could be at play between Mo&Mac and stromal or endothelial cells in FL and DLBCL, which we might have missed in this study.

## Acknowledgments

Authors wish to thank all donors and also Catherine Blanc and Aurélien Corneau, from the CyPS core facility at Sorbonne University (Paris) for access to the Helios mass cytometer; Yael Glasson from the IRCM (Montpellier) for access to the Hyperion; the H2P2 (Rennes) for slide preparation; the Biosit CytomeTri (Rennes) for cell sorting; the CRB platform (Rennes) for samples management; Steve Genebrier for CyTOF experiments; Maxime Leplat for additional graphic illustrations; Mathieu Laurichesse (Lorient) for samples preparation. Funding: Juliette Ferrant is recipient of a fellowship from the FHU CAMIn (Federation Hospitalo-Universitaire Cancer Microenvironnement et Innovation). Simon Léonard was supported by a specific grant from the LabEx IGO program (n° ANR-11-LABX-0016) funded by the «Investment into the Future» French Government program, managed by the National Research Agency (ANR). This work was supported by grants from the COmite de la REcherche Clinique et Translationnelle (CHU of Rennes), from the Ligue contre le Cancer (Comité 35 et 85), from the Institut Carnot-CALYM (Calym-Janssen 2019), from the Agence Nationale de la Recherche (ANR-17-CE15-0015 StroMAC), from the Fondation ARC pour la Recherche sur le Cancer (PGA1 RF20190208534), and from la Ligue Nationale Contre le Cancer (Equipe labellisée).

## Author contributions

Conceptualization, K.T. and M.R.; Methodology, J.F., S.L.G., F.P., K.T., and M.R; Formal analysis, J.F., S.L.G., F.P., T.P., and M.R; Investigation, J.F., S.L.G., F.P., S.L., I.P., B.H., L.D., C.M., B.A., N.V.A., C.L., H.A.M., and N.B.; Resources, C.P., A.B., V.L., and G.M.; Data curation, J.F., C.P. F.L.G., and M.R.; Writing - original draft preparation, J.F and M.R.; Writing - review and editing, J.F., S.L.G., K.T., and M.R.; Visualization, J.F., S.L.G, and M.R.; Supervision, K.T. and M.R.; Project administration, K.T. and M.R.; Funding acquisition, J.F., K.T., and M.R.

## METHODS

### KEY RESOURCES TABLE

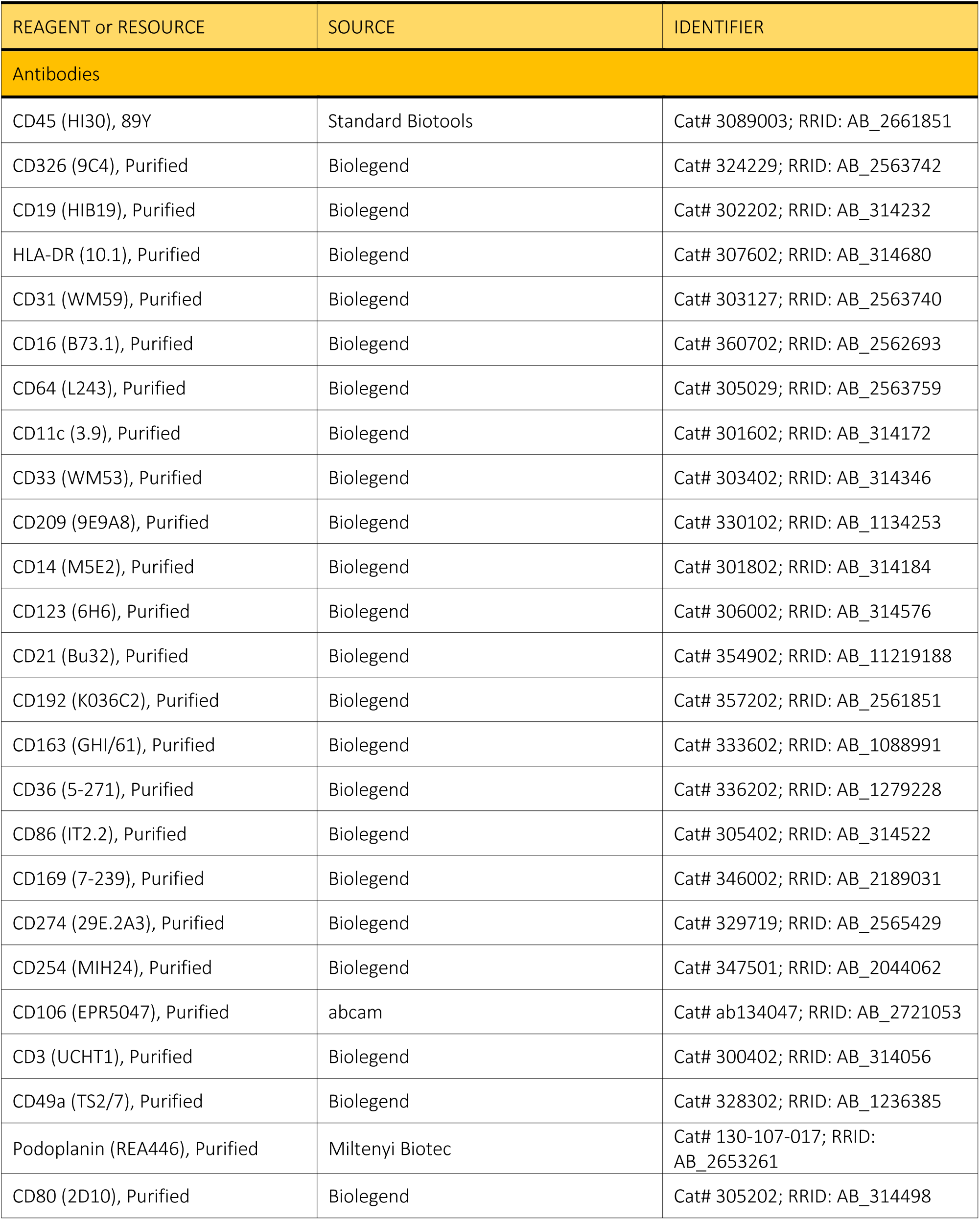
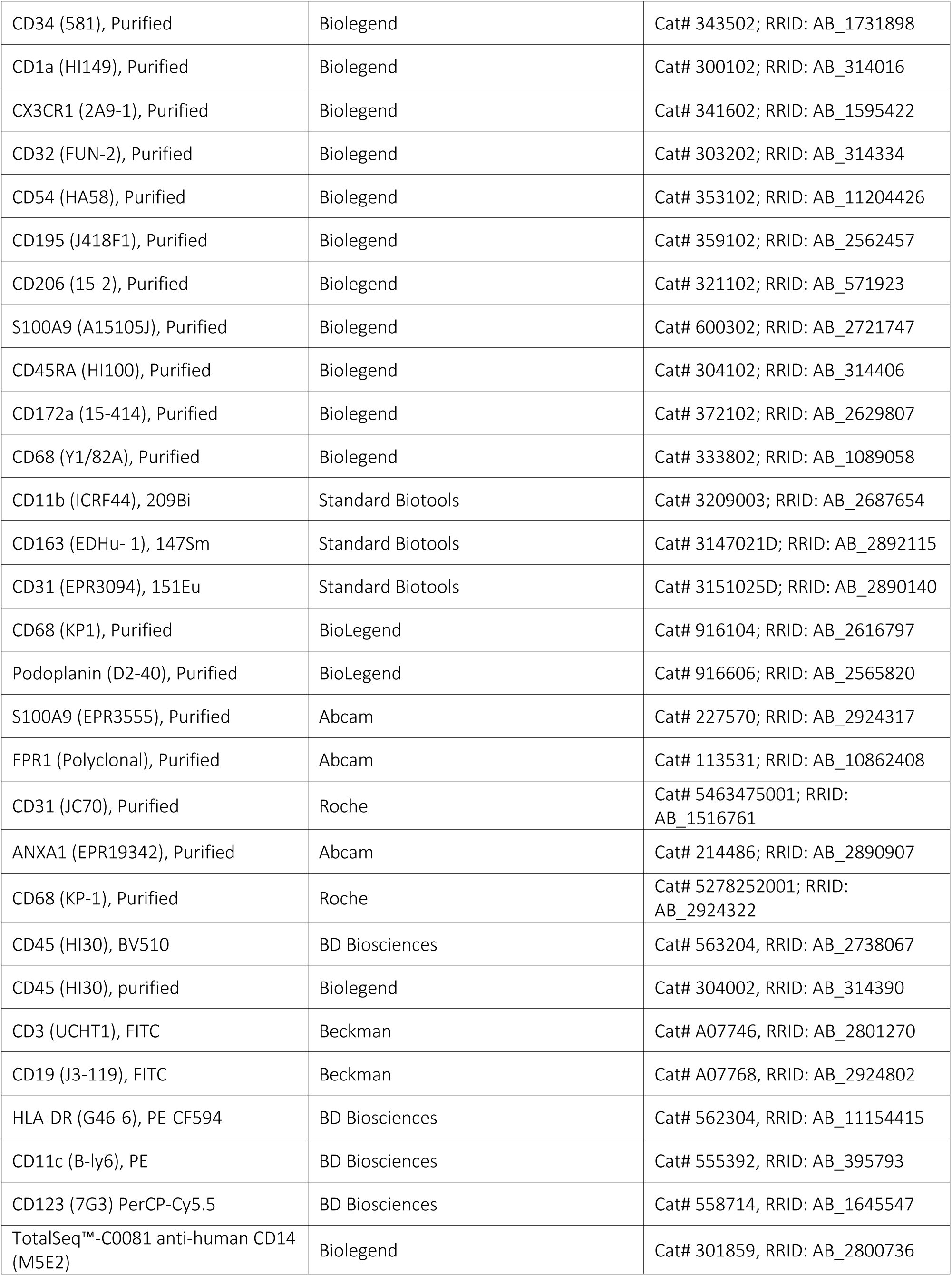
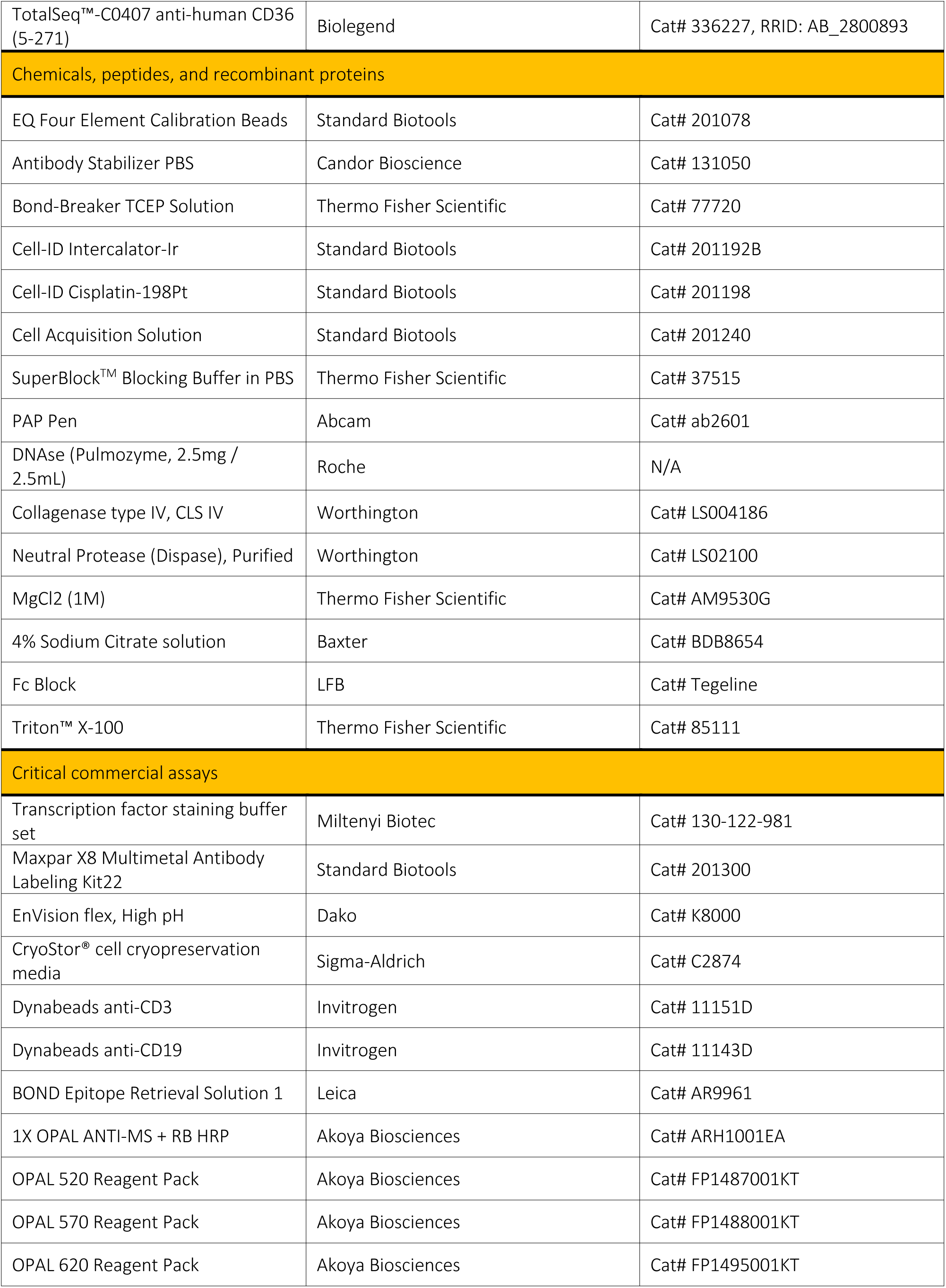
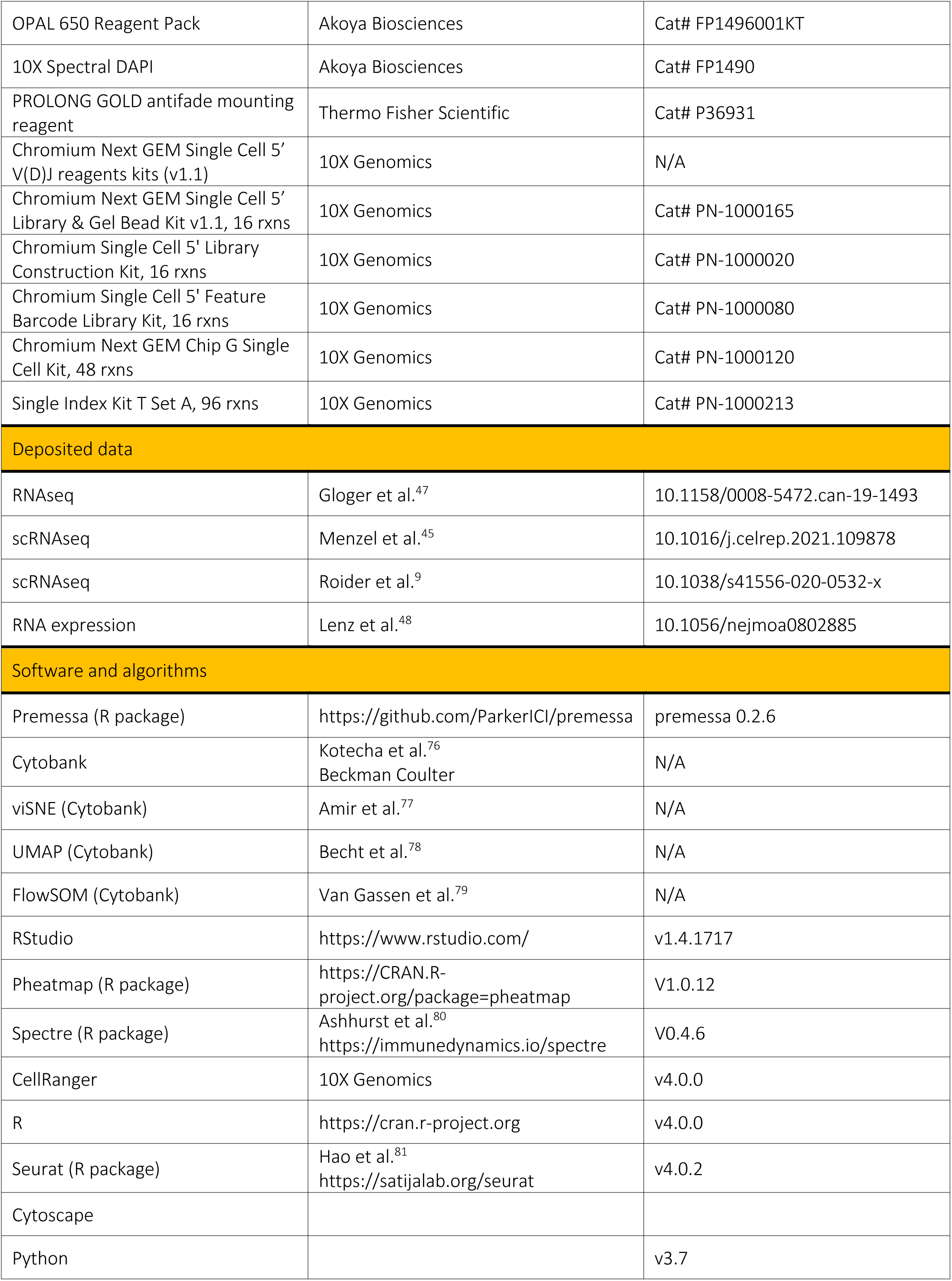
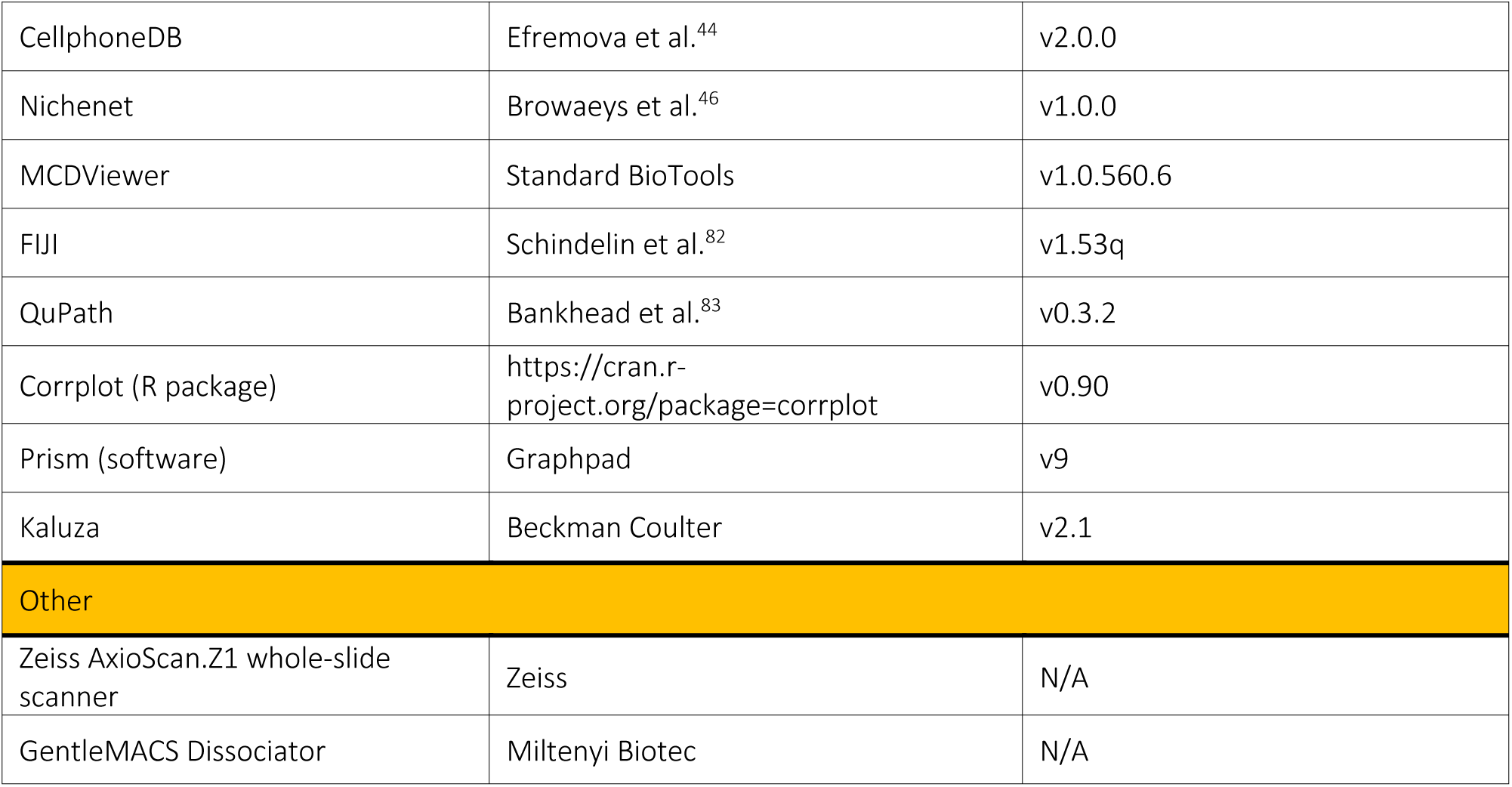

### RESOURCE AVAILABILITY

#### Lead contact

Further information and requests for resources and reagents should be directed to and will be fulfilled by the Lead Contact, Mikael Roussel

#### Material Availability

The study did not generate new unique reagents.

#### Data and Code Availability

Data generated during this study will be shared upon request.

### EXPERIMENTAL MODEL AND SUBJECT DETAILS

#### Patients and samples

Patients were recruited under institutional review board approval following informed consent process according to the declaration of Helsinki and the French National Cancer Institute ethic committee recommendations. Tissue samples included tonsils collected from routine tonsillectomy (n = 7), reactive LNs with follicular hyperplasia (n = 3), LNs from FL (n = 9) and DLBCL (n = 6) patients (Table 1). Fresh samples were minced into small fragments of about 2-3mm^3^, transferred into cryovials containing the cell cryopreservation media Cryostor CS10 (Sigma-Aldrich), and stored in liquid nitrogen. FFPE blocks from the same samples were cut into 4µm sections and placed on Superfrost Plus slides (Fisher) for *in situ* analysis.

### METHODS DETAILS

#### Tissue processing

Cell suspensions enriched in myeloid and stromal cells were obtained as previously described.^33^ Briefly, frozen tissue fragments were thawed and incubated for 5 minutes at 37°C in a dissociation buffer containing RPMI-1640 supplemented with 10 U/mL DNase (Roche), 200 U/mL collagenase IV (Worthington Biochemical), 1.6 U/mL neutral protease (Worthington Biochemical), 5 mM MgCl2, and 1% Penicillin/Streptomycin, before mechanical dissociation with a GentleMACS dissociator (Miltenyi Biotec), followed by a 40-minutes dissociation at 37°C under a 350-rpm agitation. Cell suspensions were then filtered with a 100 µm cell strainer and washed with a washed buffer containing PBS supplemented with 1% human serum albumin (HSA), 5 mM MgCl_2_, and 6.7 mg/mL sodium citrate. To block Fc receptors, cells were incubated for 5 minutes at 4°c in depletion buffer supplemented with 12.5 μg/mL Tegeline before proceeding to myeloid and stromal cells enrichment. A lymphocyte depletion using Dynabeads anti-CD3 (Invitrogen) and Dynabeads anti-CD19 (Invitrogen) was then performed. According to manufacturer’s recommendations, cells were incubated for 30 minutes at 4°C with 25 μL of Dynabeads anti-CD3 for 1 x 10^7^ cells and 25 μL of Dynabeads anti-CD19 for 2.5 x 10^7^ cells, with constant rocking on a nutating platform mixer, before magnetic removal of the beads. Cell suspensions enriched in myeloid and stromal cells were then used for CyTOF analysis or MNP FACS-sorting.

#### Antibodies and reagents for mass cytometry and imaging mass cytometry

Antibodies for mass cytometry and imaging mass cytometry were obtained already conjugated to metals (Standard BioTools) or obtained purified and then conjugated to lanthanide metals, using the MaxPar antibody conjugation kit (Standard BioTools) according to manufacturer’s recommendations.^84^ Antibodies used are listed in the Key Resources Table. Protein concentration was then determined by measurement of absorbance at 280 nm before titration on positive controls. Finally, the metal-labeled antibodies were diluted in Candor PBS Antibody Stabilization solution (Candor Bioscience) for long-term storage at 4^°^C.

#### Mass cytometry analysis

Enriched cell suspensions were prepared for CyTOF analysis as previously described.^33,84^ Briefly, cells were stained 5 minutes in RPMI 1640 supplemented with 0.5 µM Cisplatin Cell-ID (Standard BioTools) before washing with 10% FCS in RPMI 1640. Cell pellets were resuspended in 40 µl of 0.5% BSA in PBS. Surface (SB) and intracellular (IB) staining buffer were prepared with antibodies listed in the Key Resources Table. Then 60 µl of SB was added to 40 µl of resuspended cells. After staining, cells were washed in 0.5% BSA in PBS before fixation/permeabilization with a permeabilization buffer (PB) (Transcription factor staining buffer set, Miltenyi Biotec). After washing, 60µl of IB was added to 40µl of resuspended cells in PB. After intracellular staining, cells were washed twice before staining in DNA intercalator solution (2.5% Paraformaldehyde, 1:3200 Cell-ID Intercalator-Ir [Standard BioTools] in PBS). Samples were cryopreserved at −80°C until acquisition on Helios (Standard BioTools).

#### Multiplex immunofluorescence assay

The BOND RX (Leica Biosystems) was used to automate the staining procedure. After baking and dewaxing, the tissue slides were heat pre-treated using a ER1 (pH6) target retrieval buffer for 15 minutes at 97°C (AR9961, Leica Biosystems). The slides were then stained for multiplex immunofluorescence using the BOND RX Opal multiplex procedure (v7.0.3.216 RX) in a 4-step protocol with sequential denaturation (ER1 retrieval buffer (pH6), at 97 °C, AR9661, Leica Biosystems) after each staining cycle. Following endogenous peroxidase blocking (15 minutes, ambient temperature, Discovery Inhibitor, Roche, Ventana Medical Systems), tissue slides were incubated using the primary antibodies FPR1 (polyclonal, AB113531, Abcam) at a 1/250 dilution (in Envision Flex diluent (K800621-2, Agilent Technologies), CD31 (ready-to-use, clone JC70, 05463475001, Roche Diagnostics), Annexin A1 (clone EPR19342, AB214486, Abcam) at a 1/600 dilution (in Envision Flex diluent (K800621-2, Agilent Technologies), and CD68 (ready-to-use, clone KP-1, 05278252001, Roche Diagnostics). Each target was linked using the OPAL Polymer HRP conjugated secondary antibodies (ready-to-use, ARH1001EA, AKOYA Biosciences). Visualization of the different targets was established using the OPAL Polaris 570 (FP1488001KT, AKOYA Biosciences), OPAL Polaris 520 (FP1487001KT, AKOYA Biosciences), OPAL Polaris 620 (FP1495001KT, AKOYA Biosciences) and OPAL Polaris 650 (FP1496001KT, AKOYA Biosciences) detection kits, respectively. The tissue slides were counterstained using Spectral DAPI (FP1490, AKOYA Biosciences) and mounted with PROLONG GOLD antifade mounting reagent (P36931, Thermofisher). Fluorescent stained whole tissue slides were scanned in 16 bits using the Zeiss AxioScan.Z1 (Carl Zeiss) whole-slide scanner equipped with a Colibri 7 solid-state light source and appropriate filter cubes.

#### Imaging Mass Cytometry Acquisition

Two successive tissue sections were used. The first one was colored with hematoxylin-eosin-saffron (HES) to select the regions of interest to ablate. The second section was used for IMC staining. After baking and dewaxing, the tissue sections for IMC were heat pre-treated using a target retrieval solution for 30 min at 95°C (K8000, EnVision flex target retrieval solution, pH9 (Agilent Technologies). Slides were cooled down for 20 min at room temperature before rinsing with water and PBS, and were then incubated 45 min with SuperBlock Blocking Buffer (#37515, ThermoScientific). Tissue sections were next incubated overnight with antibody cocktail diluted in 0,5% BSA in PBS in humidified chamber. Antibodies used are listed in the Key Resources Table. The following day, slides were successively rinsed with 0,2% Triton X-100 in PBS (#85111, ThermoScientific) and PBS, before Iridium intercalator staining during 30 min (#201192A, Standard BioTools) and drying. Two ROI per sample were acquired using the Hyperion Imaging Mass Cytometry (Standard BioTools) after tuning the instrument according to the manufacter’s instructions. Tissues were ablated at 200 Hz.

#### MNP isolation, HTO barcoding, and CITE-seq

Cell suspensions enriched in myeloid cells were stained for 30 minutes at 4°C with different antibodies for cell sorting, cell hashing, and cellular indexing of epitopes by sequencing. Antibodies used are listed in the Key Resources Table. Cell sorting was performed on a Aria Fusion sorter. For each sample, CD45^pos^ CD3^neg^ CD19^neg^ HLA-DR^pos^ CD11c^pos^ and CD45^pos^ CD3^neg^ CD19^neg^ HLA-DR^pos^ CD11c^neg^ CD123^neg^ MNP populations were sorted and pooled, respecting their initial ratio (Figure S3A). Most samples were barcoded, we used homemade hashtag-oligo (HTO) for cell hashing, where oligo-tagged antibodies against CD45 uniquely label cells from distinct samples, which can be subsequently pooled. The homemade HTO were generated following the recommendations of the NYGCtech lab in their CITE-seq Hyper Antibody-Oligo Conjugation protocol version 2019-03-20. Samples were pooled after sorting and before library preparation, at a 1:1 cell ratio. Cytometry data are presented using Kaluza Analysis Software (Beckman Coulter).

#### Library preparation and sequencing

Sorted MNP were converted to barcoded scRNASeq gene expression and cell surface protein libraries using Chromium Next GEM Single Cell 5’ V(D)J reagents kits (v1.1) (10X Genomics) with feature barcode technology for Cell surface protein compatible with CITE-seq antibodies, according to manufacturer’s recommendations (CG000208 Rev E), with an aim of 6000 cells per library. HTO libraries were generated by PCR on 5 ng of DNA amplification product (purified for cell surface protein library) using Kapa Hifi PCR Master Mix, 10 μM 10X Genomics SI-PCR primer and 10 μM Illumina TruSeq DNA D7xx primers, followed by 1.6X SPRIselect (Beckman Coulter) purification. Library quality controls and quantifications were performed using a 2100 Bioanalyzer High Sensitivity DNA kit (Agilent). Libraries were then pooled (85% gene expression, 10% cell surface protein, 5% HTO) and subsequently sequenced on a NovaSeq 6000.

### QUANTIFICATION AND STATISTICAL ANALYSIS

#### Mass Cytometry Preprocessing

After acquisition, intrafile signal drift was normalized and .fcs files were obtained using CyTOF software. To diminish batch effects, all files were normalized on EQ Beads (Fluidigm Sciences) using the premessa R package. Files were then uploaded to the Cytobank cloud-based platform (Beckman Coulter). Data were first arcsinh-transformed using a cofactor of 5. For all files, live single cells were selected by applying a gate on DNA1 vs. DNA2 followed by a gate on DNA1 vs. Cisplatin, then beads were removed by applying a gate on the beads channel (Ce140Di) vs. DNA.1. For each sample, we performed successive viSNE, which is based upon the Barnes–Hut implementation of t-SNE, using all markers included in our panel, to remove contaminating B, T and low-quality cells, and selected all CD45^neg^ cells on one hand and all CD45^pos^ HLA-DR^pos^ CD3^neg^ CD19^neg^ cells on the other hand.

#### Mass Cytometry dimension reduction and clustering

For each population of interest, i.e. stromal/endothelial CD45^neg^ cells (594,946 cells [n = 79,955 for DLBCL, n = 105,344 for FL, n = 86,839 for rLN, and n = 322,808 for rTonsil]) and myeloid CD45^pos^ HLA-DR^pos^ CD3^neg^ CD19^neg^ cells (658,206 MNP [n = 138,374 for DLBCL, n = 152,000 for FL, n = 236,902 for rLN, and n = 130,930 for rTonsil]), we performed separate UMAP dimension reductions, using all parameters except CD3, CD19, and CD45. Then we applied a clustering method using the FlowSOM clustering algorithm. FlowSOM uses Self-Organizing Maps (SOMs) to partition cells into clusters based on their phenotype, and then builds a Minimal Spanning Tree (MST) to connect the nodes of the SOM, allowing the identification of metaclusters (i.e. group of clusters). We performed the FlowSOM algorithm on the previous UMAP results, using all events and all channels except CD3, CD19, and CD45, and the following parameters: clustering method = hierarchical consensus, iterations = 10, number of clusters = 100, number of metaclusters = 12 (for the stromal/endothelial population) or 13 (for myeloid cells). For both populations, each metacluster was manually annotated based on his marker expression phenotype, his projection on the UMAP, and his localization in the FlowSOM MST. Two stromal/endothelial metaclusters were merged into one, leading to the identification of 11 metaclusters. Single cell, clusters and metaclusters informations, including their abundances and mean marker intensities, were then exported from Cytobank for further analyses. Data representation, hierarchical clustering, and heatmaps were generated with R v4.0.0, using Rstudio v1.4.1717, and pheatmap and Spectre^80^ packages.

#### Single cell RNASeq Preprocessing

After sequencing, libraries were mapped to the human GRCh38 genome and demultiplexed using CellRanger (10X Genomics, v.4.0.0). Data from each library were then further processed using the R package Seurat (v.4.0.2) on R v.4.0.0 with Rstudio v1.4.1717. Genes expressed in three or fewer cells were excluded from downstream analysis. For each library, cells expressing a number of genes above or below 2 median absolute deviations (MAD) or a percentage of mitochondrial transcripts above 2 MAD were filtered out. Cell surface protein and HTO counts were normalized using the centered log ratio transformation (CLR) method from the NormalizeData function. HTO-tagged samples were demultiplexed using the HTODemux function with a positive quantile threshold set to 0.95. Gene expression counts were normalized using the NormalizeData function with the default scale factor of 10,000.

#### Single cell RNASeq integration, dimension reduction, and clustering

We used the standard anchor-based workflow for dataset integration in Seurat to adjust for technical and biological sources of variation between samples. All samples were then merged together and integrated data were scaled and centered using the ScaleData function. We then performed a principal component analysis (PCA) based on the top 2000 variable features followed by a graph-based clustering using the FindNeighbors and the FindCluster functions, with a resolution of 0.4. Results were visualized using a nonlinear dimensional reduction UMAP and cell clusters were annotated based on the expression of canonical markers. Then, we focused on MNP clusters. All cells included in the MNP clusters previously identified (i.e. moMac, Mac-C1Q, SIGLEC6-cDC, CLEC9A-cDC1, LAMP3-cDC, cDC2, MNDA-cDC2) were extracted, and the same integration, scaling, clustering (resolution of 0.65), annotation, and visualization workflow was applied.

#### Differentially expressed genes analysis

Differentially expressed (DE) genes analysis was performed using the FindMarker Seurat function, with a minimal fold-change of 1.5, a minimum of 30% of gene-expressing cells in at least one of the sets of cells compared, and the default Wilcoxon rank sum test. Adjusted p-values were obtained with a Bonferroni correction using all genes in the dataset. To address inter-samples heterogeneity, we performed multiple DE genes analyses while iteratively removing one of the samples from the analyses, and kept only “robust” DE genes that were found statistically DE in all sub-analyses. We performed DE gebes analyses to compare sample types (i.e. rTonsil, rLN, FL, DLBCL) for global monocyte/macrophages population as a whole (Mo&Mac metacluster) (Table S1).and then for all individual MNP clusters (i.e. CD14-moMac, S100-moMac, CD16-moMac, IL1b-moMac, Mac-C1Q) and DC clusters (i.e. MNDA-cDC2, CD1E-cDC2, CREM-cDC2, LAMP3-cDC, SIGLEC6-cDC, CLEC9A-cDC1) (Table S2).

Volcano plots were created with the R package EnhancedVolcano. Pathway enrichment analysis and visualization of DE gene was performed following Reimand et al. recommendations^42^, using g:Profiler and EnrichmentMap application on Cytoscape. DE gene lists were defined as described above and ranked by log_2_FC. Pathway enrichment analysis with g:Profiler was performed on ordered gene lists on Reactome and GO:Biological processes databases while removing not manually reviewed annotations, with a q-value <0.01, a size of query/term intersection ≥3, and a size of functional category between 5 and 350. Results were exported and subsequently uploaded in Cytoscape for visualization with EnrichmentMap, with a FDR q-value cutoff set to 1 and selection of the “Filter genes by expression” option. The prefuse force directed layout algorithm weighted using the gene set similarity coefficient was chosen for visualization. Auto annotation followed by manual arrangement of network nodes and updating of theme labels was then conducted, and nodes were colored based on their gene list of origin. Genes co-expressions were assessed using the FeatureScatter tool from the Seurat package and Pearson correlations.

#### Protein activity analysis

The R package PISCES is a regulatory-network-based methodology for the analysis of single cell gene expression profiles that allows protein activity inference.^85^ Data were normalized using SCTransform and CorDist functions, then cluster-specific metacells were generated with a minimum of 200 cells included. These metacells were used as inputs for gene networks generation with the R package corto,^86^ using as centroids DoRotheA regulons (confidence level A to E)^87^ and a p-value <1.10^-10^. After creation of a PISCES object, CPM normalization, and generation of a gene expression signature, the protein activity signature was obtained with VIPER. As protein activity includes scaled negative values, we applied a linear correction to the inferred protein data and set the minimum protein activity value to zero before DE analysis. DE analysis, volcano plots, and pathway enrichment analysis and visualization were performed as previously described for gene expression data.

#### Analysis of public datasets

We used 2 published stromal/endothelial murine lymphoma datasets for the analysis of interactions between our MNP scRNASeq dataset and BEC.^45,47^ For the CellphoneDB analysis, we collected scRNAseq data generated from Eµ-*Myc* lymphoma mice LN BEC.^45^ Using the homologene R package, we generated a conversion table of the mouse gene names into human gene names, then filtered each mouse name to match a unique human gene and created a Seurat object by matching each count to each converted mouse gene name. For the Nichenet analysis, we used bulk RNAseq data generated from the same mouse model,^47^ after a similar conversion of mouse gene names into human gene names using the homologene R package.

#### Interactome analysis

Interactome analyses were performed with CellPhoneDB (v2.0.0)^44^ on Python v3.7 and Nichenet (v1.0.0)^46^ on R v.4.0.0 with Rstudio v1.4.1717. CellPhoneDB is a manual curated repository of ligands, receptors and their interactions, integrated with a statistical framework for inferring cell-cell communication networks from single cell transcriptome data. Only receptors and ligands expressed in more than 10% of the cells in any given population were considered. We performed pairwise comparisons between all moMac clusters (Mo&Mac metacluster and all individual MNP clusters [i.e. CD14-moMac, S100-moMac, CD16-moMac, IL1b-moMac, Mac-C1Q]) from our dataset and published BEC populations from lymphoma mice.^45^ Ligand-receptor interaction scores were computed and tested using CellPhoneDB statistical analysis method with the default parameters and a p-value threshold set at 0.05. We then focused on statistically relevant predicted interactions involving up- or down-regulated genes in DLBCL vs rLN moMac, either at the RNA level or inferred with PISCES.

We used the Nichenet R package to identify top ligand-receptor and ligand-targets interactions between DLBCL-moMac and lymphoma-BEC. For this approach, we created a data matrix using the Seurat function AverageExpression on moMac from each sample to mimic bulk RNASeq data. We defined a DLBCL-moMac signature as the genes up- or down-regulated in DLBCL compared to rLN either at the RNA level or inferred with PISCES. For BEC, we used the published dataset GSE126033,^47^ after converting murine genes into human genes. Expressed genes were defined in sender and receiver cells as an average log2 expression above 0.2 for moMac and 7 for BEC. The DLBCL-moMac signature was used as the gene set of interest in the receiver population (i.e. DLBCL-moMac) to predict how ligands can impact target genes and rank receptors according to their activity. Then the best upstream ligands expressed by sender cells (i.e. BEC) were filtered.

#### Segmentation and data analysis for imaging mass cytometry

MCD files were converted to ome.tiff format through MCDViewer (Standard BioTools). We normalized all ROI (99th percentile normalization) and performed a scaling between 0 to 100 using FIJI software.^82^ Blood vessels were constituted of BEC defined as CD31^pos^ PDPN^neg^ annotations using the QuPath Software’s pixel classification.^83^ Areas of 25µm around the blood vessels structure were arbitrary defined using the Expand annotations function in QuPath. Due to their complex morphology, macrophages were segmented using the cell detection command in QuPath based on the expression of CD68, CD163, and S100A9. CSV files of cells detections were exported from Qupath for further analyses.

#### Statistical analysis

Statistical analyses were performed with Graphpad Prism 9.0.0 or R v4.0.0, using Rstudio v1.4.1717. P values were defined by a Kruskal-Wallis test followed by a Dunn’s post-test for multiple group comparisons or by Wilcoxon matched-pairs signed rank tests as appropriate. Correlations were calculated using Spearman test. * p < 0.05, ** p < 0.01, *** < 0.001. Heatmaps were generated with the pheatmap package and correlation matrixes were built with the corrplot package. Hierarchical clustering of the patients was performed using euclidean distance and ward.D2 clustering.

