## Supplemental Figures for "Single-cell profiling identifies clinically relevant interactions between tumor associated macrophages and blood endothelial cells in diffuse large B cell lymphoma"

### Figure Legend

#### Figure S1: CyTOF panel and analysis pipeline (Related to Figure 1)

**(A)** Experimental design overview.

**(B)** CyTOF panel dedicated to the study of SLO MNP and stromal/endothelial cells.

**(C)** CyTOF analysis flowchart.

**(D)** Abundances of significantly enriched MNP clusters in lymphoma, according to sample type. Mann-Whitney, *p<0.05, **p<0.01.

#### Figure S2: Myeloid, stroma, and endothelial phenotype (Related to Figure 2)

(A) Heatmap of relative proportions of all CyTOF clusters and relevant mean metal intensity (MMI) in distinct clusters between sample types (FL, DLBCL, rLN, rTonsil). Statistically different parameters are highlighted. Statistically different relative proportions and relevant statistically different MMI (i.e. number of cells in compared clusters > 500, arcsinh transformed MMI > 1 in at least one condition for the compared clusters, and statistical difference not only relying on rTonsil tissues) were selected for correlation analyses (See Figure 2). Kruskal-Wallis, p<0.05.

(B) Boxplots of selected CyTOF clusters abundances and selected markers MMI for chosen differentially expressed clusters, according to sample type. Kruskal-Wallis, multiple comparison with Dunn correction, *p<0.05, **p<0.01, ***p<0.001.

#### Figure S3: Cell sorting and scRNAseq flowchart (Related to Figure 3)

(A) Gating strategy to sort OLS MNP – excluding pDC – by FACS.

(B) scRNASeq analysis flowchart.

(C) UMAP and FlowSOM clustering of all viable cells from SLO included in the single cell RNASeq analysis, displaying the 17 clusters identified, including 7 non-pDC MNP clusters (top left), or the cells colored by sample – i.e. DLBCL, FL, rLN, rTonsil (bottom left), and projection of selected markers expression (right).

(D) Boxplots of scRNASeq clusters abundances according to sample type. Kruskal-Wallis, multiple comparison with Dunn correction.

**Figure S4: Monocytes and macrophages transcriptomic deregulation in lymphoma (Related to Figure 4)**

(A) Principal component analysis (PCA) on top 1000 expressed genes in rLN and rTonsil samples, barycenter of each group is shown. Volcano plots of differentially expressed genes for all cells (i.e. monocytes/macrophages) included in the 5 monocytes/macrophages clusters – CD14-moMac, S100-moMac, CD16-moMac, IL1b_moMac/DC, Mac-C1Q – between rTonsil and rLN (left), rTonsil and FL (center), rTonsil and DLBCL (right). Wilcoxon-Mann-Whitney test, $|FC| > 1.5$, $p\text{-val}_{adj}<0.05$. Only genes present in at least 30% of the cells in one of the compared conditions and found to be robustly differentially expressed by iteratively removing each sample from the differential analysis are presented as differentially expressed (larger, colored dots).

(B) Volcano plots of statistically different inferred proteins for all cells (i.e. monocytes/macrophages) included in the 5 monocytes/macrophages clusters – CD14-moMac, S100-moMac, CD16-moMac, IL1b_moMac/DC, Mac-C1Q – between FL and DLBCL (left), DLBCL and reactive LN (center), FL and reactive LN (right). Wilcoxon-Mann-Whitney test, $|FC| > 1.5$, $p<0.05$. Only proteins present in at least 30% of the cells in one of the compared conditions and found to be robustly differentially expressed by iteratively removing each sample from the differential analysis are presented as differentially expressed (larger, colored dots). Proteins found to be deregulated in the comparison between clusters of one sample type and the other two sample types are highlighted in darker colors. Proteins were inferred using the PISCES method.[1]

(C) Enrichment map of the pathway analysis for the differentially expressed inferred proteins. The pathway analysis was conducted with g:profiler, using the Reactome and GO:BP databases and the enrichment map was built in Cytoscape by combining all 6 pathway analyses corresponding to all comparisons between FL, DLBCL and rLN samples.[2] Each node represents a pathway, colored according to the context in which it is enriched. Pathways were found enriched only in the DLBCL vs FL and DLBCL vs rLN signatures.

**Figure S5: Interactome and *in situ* analyses identifies BEC and Mo&Mac crosstalk in DLBCL (Related to Figure 5)**

(A) Dotplot of statistically significant interactions identified with CellphoneDB[3] between normal murine BEC and rLN moMac clusters or tumor murine BEC and DLBCL moMac clusters, involving ligands expressed by murine BEC and receptors expressed by moMac and differentially expressed between rLN and DLBCL LN. Log2 mean expression of interacting molecules (color scale) and p-values (dot size) are shown. Only statistically significant interactions are plotted ($p<0.05$) with a minimal p-value set to 0.01.

(B) Dotplot of statistically significant interactions identified with CellphoneDB[3] between normal murine BEC and rLN moMac or tumor murine BEC and DLBCL moMac, involving ligands expressed by moMac and receptors expressed by murine BEC and differentially expressed between rLN and DLBCL LN. Log2 mean expression of interacting molecules (color scale) and p-values (dot size) are shown. Only statistically significant interactions are plotted ($p<0.05$) with a minimal p-value set to 0.01 (left). Dotplots showing the average RNA expression level (color scale) and percentage of expressing cells (dot size) of the identified ligands expressed by moMac and receptors expressed by tumor or normal BEC (right).

(C) Heatmap of the interaction potential between the top 12 ligands expressed by tumor murine BEC identified with Nichenet and their receptors expressed by DLBCL moMac. Dotplots showing the average RNA expression level (color scale) and percentage of expressing cells (dot size) of the identified ligands expressed by tumor BEC and receptors expressed by Mo&Mac.

(D) Boxplots showing ANXA1, FPR1 and FPR2 expression levels on BEC, LEC, FRC and DN from naïve or tumor mouse LN tissue (GSE126033).[4]

(E) Violin plots showing ANXA1, FPR1 and FPR2 expression levels on T and B cells from DLBCL LN and rLN.[5]

(F) Violin plot showing ANXA1 expression level on DLBCL and rLN Mo&Mac.

(G) CD31, ANXA1, CD68, and FPR1 staining by mIF in DLBCL and rLN. Scale bar represent 20 µm.

(H) Top 10 genes correlated to FPR1 or FPR2 in DLBCL moMac. Pearson correlation ***=$p<0.001$.

**Figure S6: *In situ* analyses identifies BEC and Mo&Mac crosstalk in DLBCL (Related to Figure 5)**

IMC staining for BEC ($CD31^{pos}$ $PDPN^{neg}$), LEC ($CD31^{pos}$ $PDPN^{pos}$), and macrophages/myeloid cells (CD68, CD163, and S100A9) in whole ROI. ROI for DLBCL (n=4, 10 ROI, total area 7.1 $mm^2$), FL (n=3,

5 ROI, total area 5.6 $mm^2$), and rLN (n=3, 6 ROI, total area 6.5 $mm^2$). Blood vessels were defined as anatomical structure $CD31^{pos}$ $PDPN^{neg}$. CD31 cells are in yellow; PDPN are in red, CD68 are in purple, CD163 are in green, and S100A9 are in pink; neighbourhoods' zones of 25 µm around blood vessels are indicated in light green. Representative ROI for DLBCL, FL, and rLN.

Blood vessel, number of macrophages, and macrophages expressing CD163 and/or S100A9 in all ROI. Kruskal-Wallis with Dunn's multiple comparisons correction, *$p < 0.05$. Scale bar represent 100 µm.


1. Obradovic, A., Vlahos, L., Laise, P., Worley, J., Tan, X., Wang, A., and Califano, A. (2022). PISCES: A pipeline for the Systematic, Protein Activity-based Analysis of Single Cell RNA Sequencing Data. Biorxiv, 2021.05.20.445002. 10.1101/2021.05.20.445002.

2. Reimand, J., Isserlin, R., Voisin, V., Kucera, M., Tannus-Lopes, C., Rostamianfar, A., Wadi, L., Meyer, M., Wong, J., Xu, C., et al. (2019). Pathway enrichment analysis and visualization of omics data using g:Profiler, GSEA, Cytoscape and EnrichmentMap. Nat Protoc *14*, 482–517. 10.1038/s41596-018-0103-9.

3. Efremova, M., Vento-Tormo, M., Teichmann, S.A., and Vento-Tormo, R. (2020). CellPhoneDB: inferring cell–cell communication from combined expression of multi-subunit ligand–receptor complexes. Nat Protoc *15*, 1484–1506. 10.1038/s41596-020-0292-x.

4. Gloger, M., Menzel, L., Grau, M., Vion, A.-C., Anagnostopoulos, I., Zapukhlyak, M., Gerlach, K., Kammertöns, T., Hehlgans, T., Zschummel, M., et al. (2020). Lymphoma Angiogenesis Is Orchestrated by Noncanonical Signaling Pathways. Cancer Res *80*, 1316–1329. 10.1158/0008-5472.can-19-1493.

5. Roider, T., Seufert, J., Uvarovskii, A., Frauhammer, F., Bordas, M., Abedpour, N., Stolarczyk, M., Mallm, J.-P., Herbst, S.A., Bruch, P.-M., et al. (2020). Dissecting intratumour heterogeneity of nodal B-cell lymphomas at the transcriptional, genetic and drug-response levels. Nature cell biology, 1–27. 10.1038/s41556-020-0532-x.

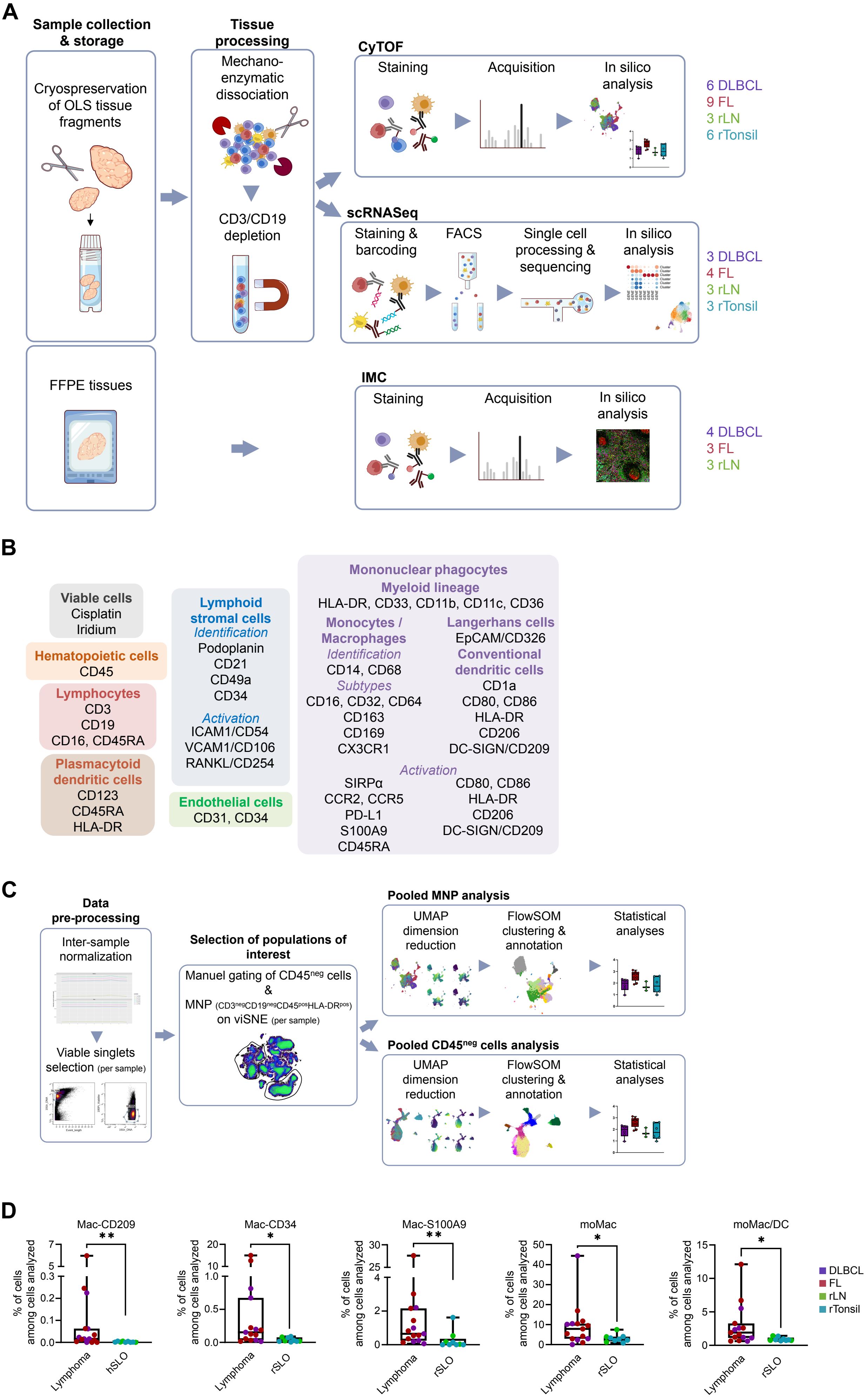
A
Sample collection & storage
Cryospreservation of OLS tissue fragments
FFPE tissues
Tissue processing
Mechano-enzymatic dissociation
CD3/CD19 depletion
CyTOF
Staining
Acquisition
In silico analysis
6 DLBCL
9 FL
3 rLN
6 rTonsil
scRNASeq
Staining & barcoding
FACS
Single cell processing & sequencing
In silico analysis
3 DLBCL
4 FL
3 rLN
3 rTonsil
IMC
Staining
Acquisition
In silico analysis
4 DLBCL
3 FL
3 rLN
B
Viable cells
Cisplatin
Iridium
Hematopoietic cells
CD45
Lymphocytes
CD3
CD19
CD16, CD45RA
Plasmacytoid dendritic cells
CD123
CD45RA
HLA-DR
Lymphoid stromal cells
Identification
Podoplanin
CD21
CD49a
CD34
Activation
ICAM1/CD54
VCAM1/CD106
RANKL/CD254
Endothelial cells
CD31, CD34
Mononuclear phagocytes
Myeloid lineage
HLA-DR, CD33, CD11b, CD11c, CD36
Monocytes / Macrophages
Identification
CD14, CD68
Subtypes
CD16, CD32, CD64
CD163
CD169
CX3CR1
Langerhans cells
EpCAM/CD326
Conventional dendritic cells
CD1a
CD80, CD86
HLA-DR
CD206
DC-SIGN/CD209
Activation
SIRPα
CCR2, CCR5
PD-L1
S100A9
CD45RA
CD80, CD86
HLA-DR
CD206
DC-SIGN/CD209
C
Data pre-processing
Inter-sample normalization
Viable singlets selection (per sample)
Selection of populations of interest
Manuel gating of CD45neg cells
&
MNP (CD3negCD19negCD45posHLA-DRpos) on viSNE (per sample)
Pooled MNP analysis
UMAP dimension reduction
FlowSOM clustering & annotation
Statistical analyses
Pooled CD45neg cells analysis
UMAP dimension reduction
FlowSOM clustering & annotation
Statistical analyses
D
Mac-CD209
Mac-CD34
Mac-S100A9
moMac
moMac/DC
% of cells among cells analyzed
Lymphoma
rSLO
DLBCL
FL
rLN
rTonsil

Figure S1

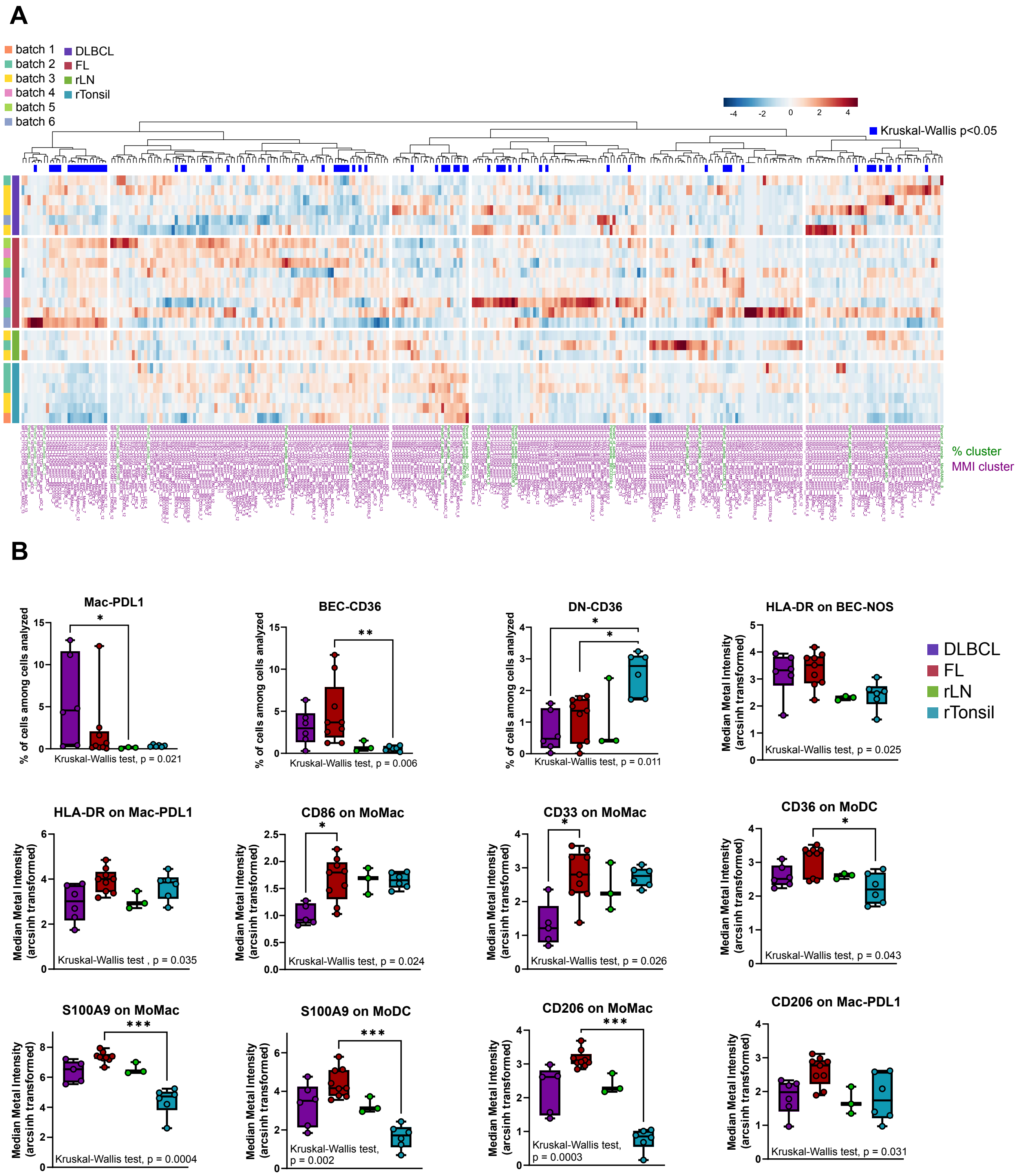
A
batch 1
batch 2
batch 3
batch 4
batch 5
batch 6
DLBCL
FL
rLN
rTonsil
-4 -2 0 2 4
Kruskal-Wallis p<0.05
% cluster
MMI cluster
B
Mac-PDL1
% of cells among cells analyzed
Kruskal-Wallis test , p = 0.021
BEC-CD36
Kruskal-Wallis test, p = 0.006
DN-CD36
Kruskal-Wallis test, p = 0.011
HLA-DR on BEC-NOS
Median Metal Intensity (arcsinh transformed)
Kruskal-Wallis test, p = 0.025
DLBCL
FL
rLN
rTonsil
HLA-DR on Mac-PDL1
Kruskal-Wallis test , p = 0.035
CD86 on MoMac
Kruskal-Wallis test, p = 0.024
CD33 on MoMac
Kruskal-Wallis test, p = 0.026
CD36 on MoDC
Kruskal-Wallis test, p = 0.043
S100A9 on MoMac
Kruskal-Wallis test, p = 0.0004
S100A9 on MoDC
Kruskal-Wallis test, p = 0.002
CD206 on MoMac
Kruskal-Wallis test, p = 0.0003
CD206 on Mac-PDL1
Kruskal-Wallis test, p = 0.031

Figure S2

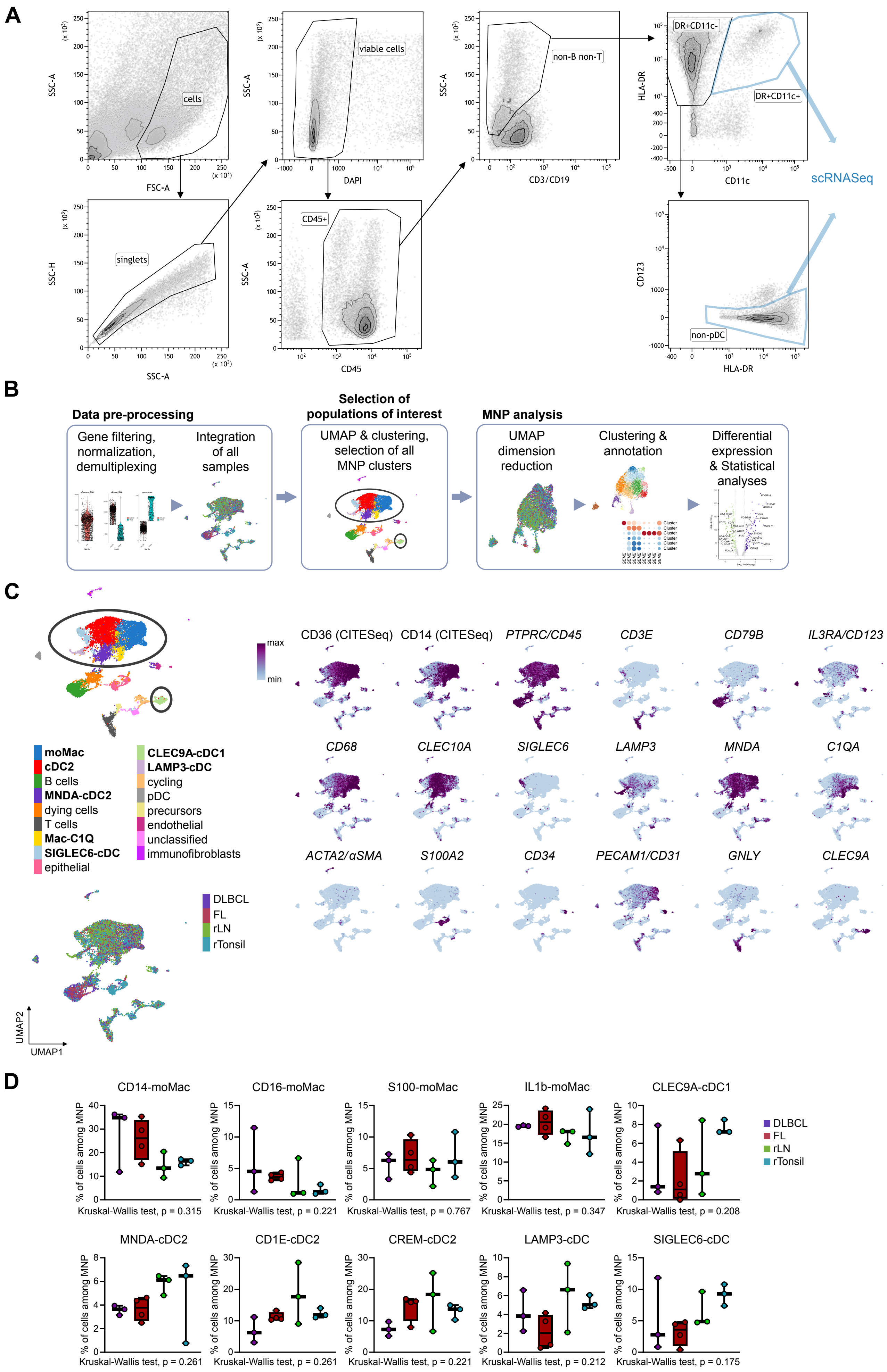
A
SSC-A
FSC-A
cells
SSC-H
singlets
DAPI
viable cells
CD45
CD45+
CD3/CD19
non-B non-T
HLA-DR
CD11c
DR+CD11c-
DR+CD11c+
CD123
non-pDC
scRNASeq
B
Data pre-processing
Gene filtering, normalization, demultiplexing
Integration of all samples
Selection of populations of interest
UMAP & clustering, selection of all MNP clusters
MNP analysis
UMAP dimension reduction
Clustering & annotation
Differential expression & Statistical analyses
C
moMac
cDC2
B cells
MNDA-cDC2
dying cells
T cells
Mac-C1Q
SIGLEC6-cDC
epithelial
CLEC9A-cDC1
LAMP3-cDC
cycling
pDC
precursors
endothelial
unclassified
immunofibroblasts
DLBCL
FL
rLN
rTonsil
UMAP2
UMAP1
max
min
CD36 (CITESeq)
CD14 (CITESeq)
PTPRC/CD45
CD3E
CD79B
IL3RA/CD123
CD68
CLEC10A
SIGLEC6
LAMP3
MNDA
C1QA
ACTA2/αSMA
S100A2
CD34
PECAM1/CD31
GNLY
CLEC9A
D
CD14-moMac
CD16-moMac
S100-moMac
IL1b-moMac
CLEC9A-cDC1
% of cells among MNP
Kruskal-Wallis test, p = 0.315
Kruskal-Wallis test, p = 0.221
Kruskal-Wallis test, p = 0.767
Kruskal-Wallis test, p = 0.347
Kruskal-Wallis test, p = 0.208
DLBCL
FL
rLN
rTonsil
MNDA-cDC2
CD1E-cDC2
CREM-cDC2
LAMP3-cDC
SIGLEC6-cDC
Kruskal-Wallis test, p = 0.261
Kruskal-Wallis test, p = 0.261
Kruskal-Wallis test, p = 0.221
Kruskal-Wallis test, p = 0.212
Kruskal-Wallis test, p = 0.175

Figure S3

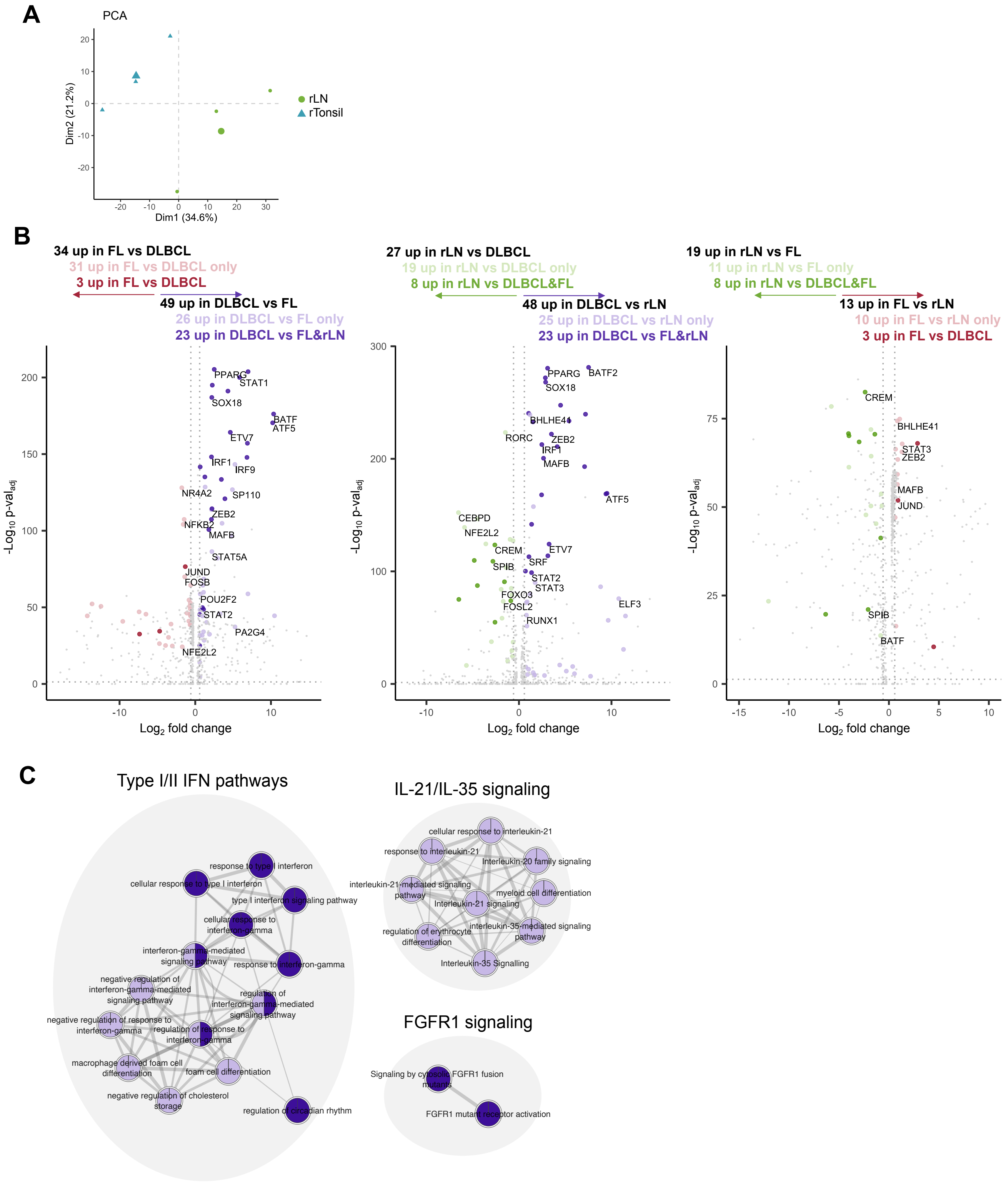

A
PCA
rLN
rTonsil
Dim1 (34.6%)
Dim2 (21.2%)
B
34 up in FL vs DLBCL
31 up in FL vs DLBCL only
3 up in FL vs DLBCL
49 up in DLBCL vs FL
26 up in DLBCL vs FL only
23 up in DLBCL vs FL&rLN
27 up in rLN vs DLBCL
19 up in rLN vs DLBCL only
8 up in rLN vs DLBCL&FL
48 up in DLBCL vs rLN
25 up in DLBCL vs rLN only
23 up in DLBCL vs FL&rLN
19 up in rLN vs FL
11 up in rLN vs FL only
8 up in rLN vs DLBCL&FL
13 up in FL vs rLN
10 up in FL vs rLN only
3 up in FL vs DLBCL
-Log10 p-valadj
Log2 fold change
C
Type I/II IFN pathways
IL-21/IL-35 signaling
FGFR1 signaling


Figure S4

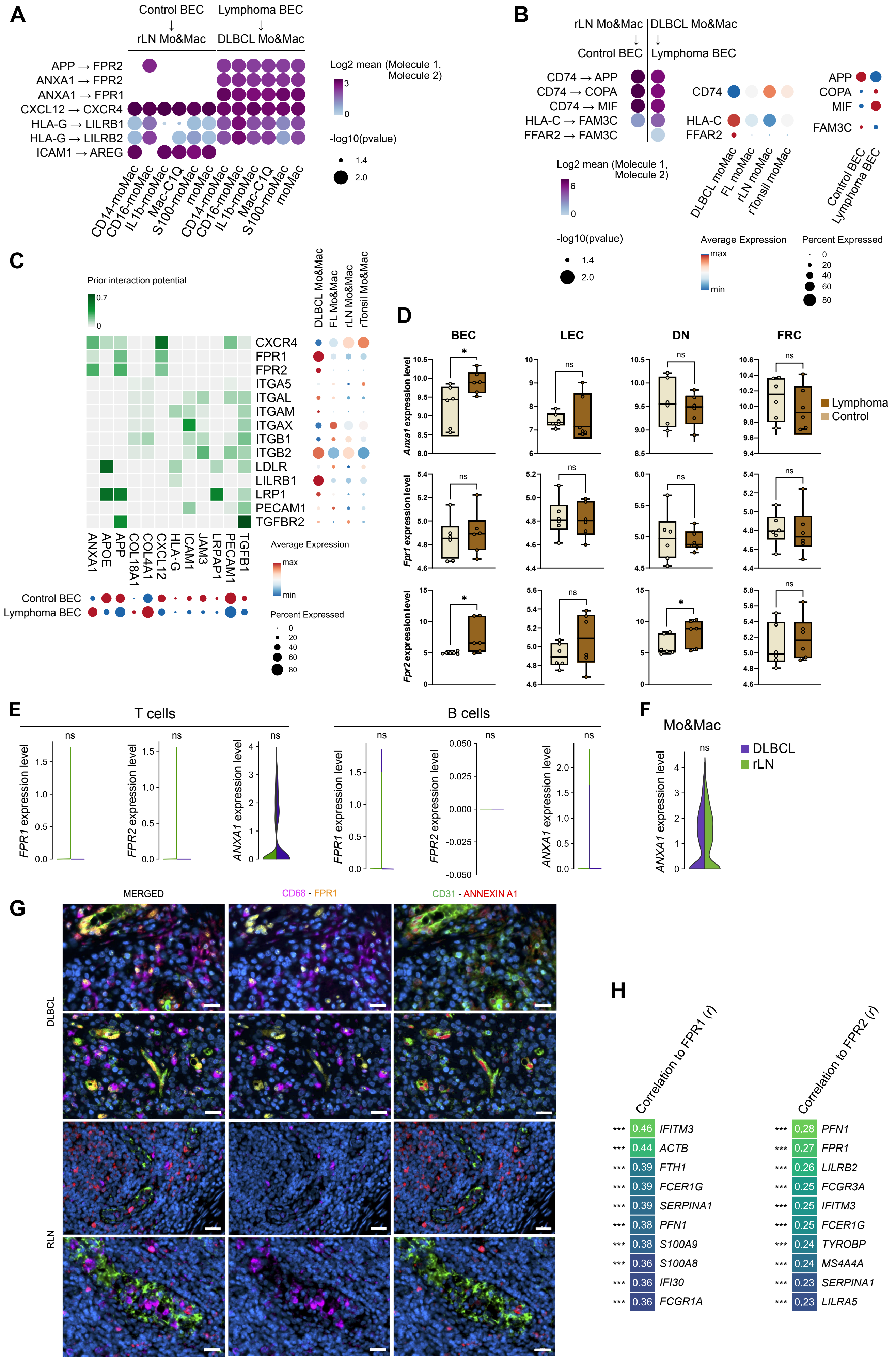
A
Control BEC → rLN Mo&Mac
Lymphoma BEC → DLBCL Mo&Mac
APP → FPR2
ANXA1 → FPR2
ANXA1 → FPR1
CXCL12 → CXCR4
HLA-G → LILRB1
HLA-G → LILRB2
ICAM1 → AREG
Log2 mean (Molecule 1, Molecule 2)
-log10(pvalue)
B
rLN Mo&Mac → Control BEC
DLBCL Mo&Mac → Lymphoma BEC
CD74 → APP
CD74 → COPA
CD74 → MIF
HLA-C → FAM3C
FFAR2 → FAM3C
Average Expression
Percent Expressed
C
Prior interaction potential
CXCR4
FPR1
FPR2
ITGA5
ITGAL
ITGAM
ITGAX
ITGB1
ITGB2
LDLR
LILRB1
LRP1
PECAM1
TGFBR2
Control BEC
Lymphoma BEC
D
BEC
LEC
DN
FRC
Anxa1 expression level
Fpr1 expression level
Fpr2 expression level
Lymphoma
Control
E
T cells
B cells
FPR1 expression level
FPR2 expression level
ANXA1 expression level
F
Mo&Mac
DLBCL
rLN
G
MERGED
CD68 - FPR1
CD31 - ANNEXIN A1
DLBCL
RLN
H
Correlation to FPR1 (r)
*** 0.46 IFITM3
*** 0.44 ACTB
*** 0.39 FTH1
*** 0.39 FCER1G
*** 0.39 SERPINA1
*** 0.38 PFN1
*** 0.38 S100A9
*** 0.36 S100A8
*** 0.36 IFI30
*** 0.36 FCGR1A
Correlation to FPR2 (r)
*** 0.28 PFN1
*** 0.27 FPR1
*** 0.26 LILRB2
*** 0.25 FCGR3A
*** 0.25 IFITM3
*** 0.25 FCER1G
*** 0.24 TYROBP
*** 0.24 MS4A4A
*** 0.23 SERPINA1
*** 0.23 LILRA5

Figure S5

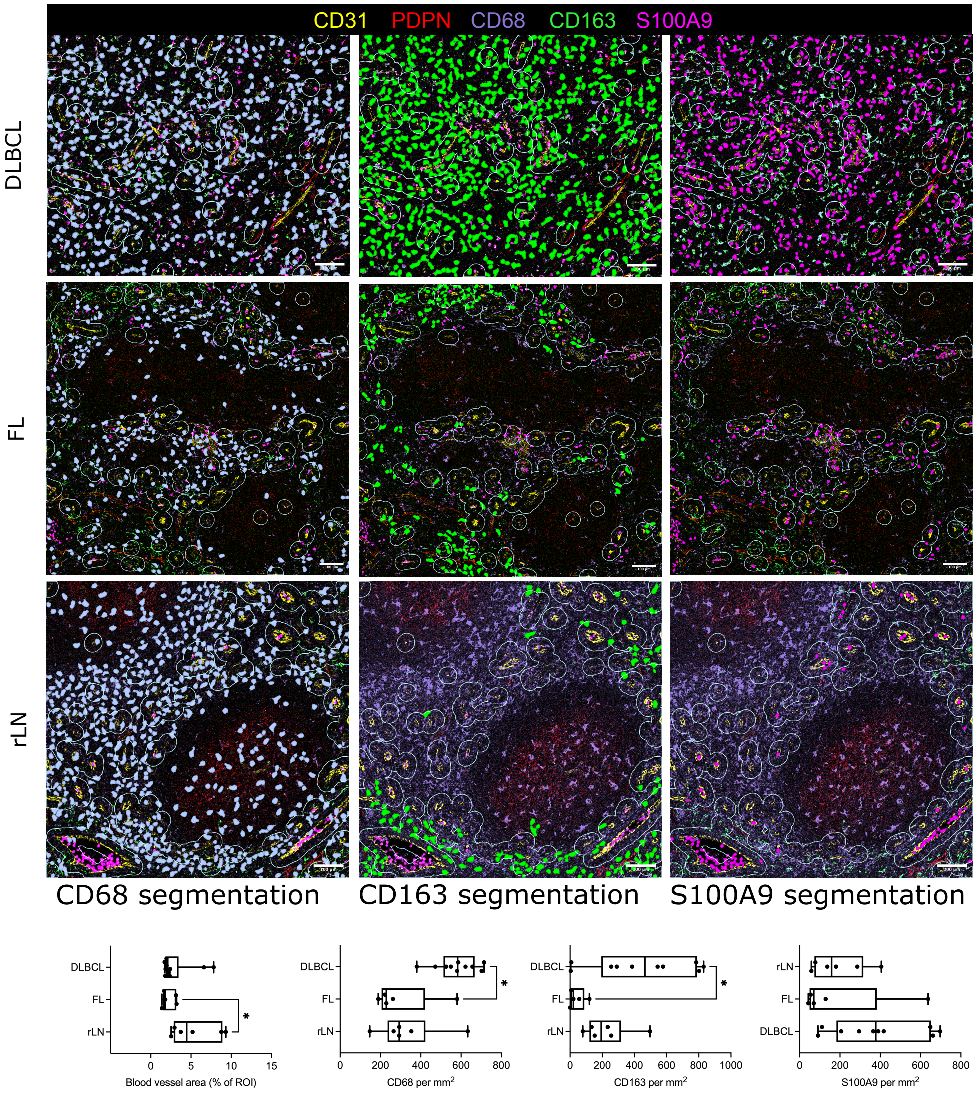
CD31 PDPN CD68 CD163 S100A9
DLBCL
FL
rLN
CD68 segmentation
CD163 segmentation
S100A9 segmentation
DLBCL
FL
rLN
Blood vessel area (% of ROI)
CD68 per mm²
CD163 per mm²
S100A9 per mm²
*

Figure S6
